# Evaluation of extraction solvents for untargeted metabolomics analysis of enrichment reactor cultures performing enhanced biological phosphorus removal (EBPR)

**DOI:** 10.1101/573535

**Authors:** Nay Min Min Thaw Saw, Pipob Suwanchaikasem, Rogelio Zuniga-Montanez, Guanglei Qiu, Stefan Wuertz, Rohan B. H. Williams

## Abstract

**Introduction:** The extraction solvent mixtures were optimized for untargeted metabolomics analysis of microbial communities from two laboratory scale activated sludge reactors performing enhanced biological phosphorus removal (EBPR).

**Objective:** To develop a robust and simple analytical protocol to analyse microbial metabolomics from EBPR bioreactors.

**Methods:** Extra- and intra-cellular metabolites were extracted using five methods and analysed by ultraperformance liquid chromatography mass spectrometry (UPLC-MS).

**Results:** The optimal extraction method was biomass specific and methanol:water (1:1 v/v) and methanol:chloroform:water (2:2:1 v/v) were chosen, respectively, for each of the two different bioreactors.

**Conclusion:** Our approach provides direct surveys of the metabolic state of PAO-enriched EBPR communities, showing that extraction methods should be carefully tailored to the microbial community under study

## Introduction

Complex microbial communities are responsible for the removal of specific pollutants in biological wastewater treatment (Cydzik-Kwiatkowska and Zielińska 2016). Despite decades of study, considerable gaps remain between the knowledge of the bioprocess engineering aspects of wastewater treatment and the underlying microbial ecophysiology of activated sludge communities, which may limit stability, optimisation and future innovation in water reclamation technologies (Cydzik-Kwiatkowska and Zielińska 2016). Recent use of omics technologies has provided many new insights, but to date most activity has been focused on using whole community metagenomics and metatranscriptomics, with very limited use of community-wide metabolomics studies (Narayanasamy et al. 2015). Notably there is little information available on optimal metabolome extraction procedures for the suspended flocular biofilms that typically represent the biomass of activated sludge communities.

Here we present a complete analytical protocol for untargeted metabolomics analysis of microbial communities from two laboratory scale activated sludge reactors designed to enrich for polyphosphate accumulating organisms (PAOs) (Kawakoshi et al. 2012). Specifically, the aim of this study is to optimize solvent conditions for sample preparation of microbial cells in activated sludge bioreactor communities that are performing enhanced biological phosphorus removal (EBPR), a key bioprocess commonly used in wastewater treatment. In two different EBPR reactor microbial communities, extra- and intra-cellular metabolites were extracted using different solvent mixtures and analyzed by ultra-performance liquid chromatography mass spectrometry (UPLC-MS), with evaluation primarily based on sample reproducibility and the breadth of metabolome coverage.

## Materials and Methods

### Operation of the lab-scale SBR enrichment bioreactors

We sampled two laboratory enrichment bioreactor communities operated using two different protocols, each designed to enrich different PAOs, from here on described as Reactor A and Reactor B (detailed methods for the operation of each reactor are described in Supplementary Methods). Enrichment reactors are commonly used in environmental engineering, to both increase the relative abundance of unculturable member species, and to provide a stable community for investigating a specific bioprocess or functional phenotype (Schlegel and Jannasch 1967).

Reactor A was a sequencing batch reactor (SBR) with a 5.4 L working volume, and was inoculated with activated sludge obtained from an existing EBPR enrichment reactor, with a protocol designed to enrich for known PAO species from the genus *Candidatus* Accumulibacter (Supplementary Material 2 - Fig 1). Analysis of community composition using 16S amplicon sequencing demonstrated the presence of around 445 distinct taxa (as captured by 16S amplicon sequences, see Hugerth and Andersson, 2017), with *Ca.* Accumulibacter accounting for approximately 27% of the biomass. (data not shown). The SBR was operated with 6 h cycles, including a feeding (60 min) stage, an anaerobic stage (20 min), an aerobic stage (180 min), and a settling/decant stage (100 min). The pH was controlled at 7.20-7.60 with dissolved oxygeon (DO) levels maintained at 0.8-1.2 mg/L during the aerobic phase.

**Fig 1.**
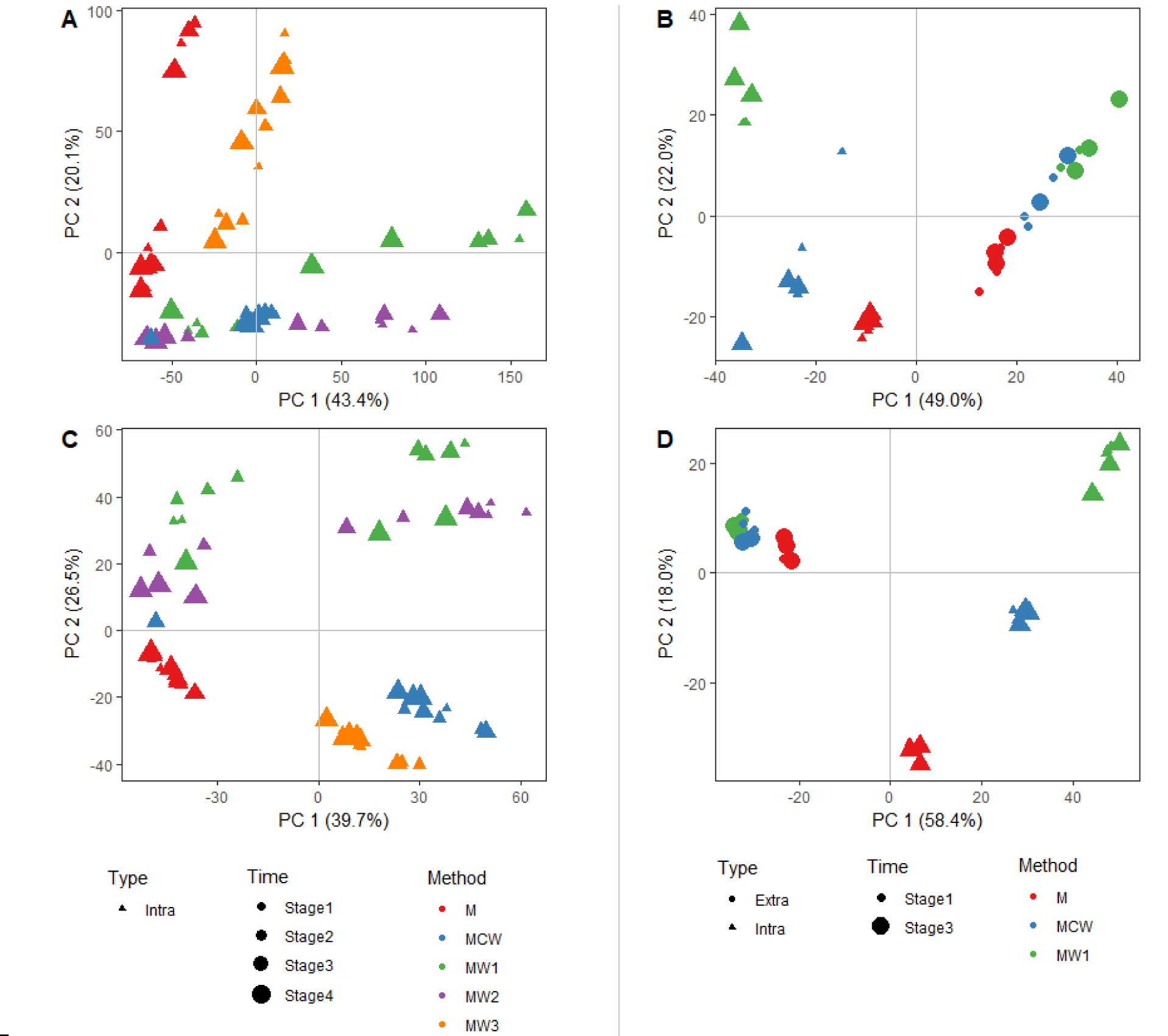
Principal components analysis of metabolite profiles of Reactor A (panel A and C) and Reactor B (B and D) extracted by different extraction methods, and categorised by ionisation mode (positive mode in panels A-B; negative mode in panels C-D)

Reactor B is a 5-L SBR containing a microbial consortium capable of performing EBPR (Supplementary Material 2 - Fig 2), and was fed with synthetic wastewater containing glucose as the main carbon source. Analysis of 16S amplicon sequencing data demonstrates the presence of around 358 taxa with the dominant member being from the genus *Nocardioides*, and constituting 23% of the biomass (data not shown). The cycle duration was 6 h and consisted of 30 min feeding, 125 min anaerobic, 154 min aerobic, 1 min sludge discharge, 35 min settling and 15 min supernatant discharge phases. The temperature was set at 31 ± 1° C and the pH was controlled at 7.5 ± 0.25 by using 0.5 M NaOH and 0.5 M HCl.

### Experimental design, sample collection and extraction procedures

We obtained samples from Reactor A and Reactor B during one 6h cycle study. For Reactor A, the sampled time points were at 30 min, 70 min, 95 min and 200 min relative to the start of the feeding phase. For Reactor B, the sampled time points were at 30 min and 95 min relative to the start of the cycle. At each time point, a 150 mL aliquot of activated sludge was sampled. Three technical replicates were obtained from each sampled aliquot. The samples were centrifuged at 10000 rpm for 1 min at 4°C. Cell pellets and supernatant were separated and snap-frozen in liquid nitrogen and stored at −80°C before lyophilization. For Reactor A, only the pellets were used to extract intracellular metabolites, while both pellet and supernatant were used to extract extra- and intracellular metabolites for the Reactor B. In order to normalize the sample volume, 10 ml of supernatant from each sample was lyophilized. After lyophilization, the samples were subjected to different metabolite extraction methods. We used up to five different extraction solutions, namely: pure methanol (M), methanol:water (50:50 v/v) (MW1), methanol:water (60:40 v/v) (MW2), methanol:water (80:20 v/v) (MW3) and methanol:chloroform:water (40:40:20 v/v/v) (MCW). For reactor A, we tested all five extraction solutions to examine the entire cellular components of intracellular metabolites, while for reactor B, M, MW1 and MCW were applied to extract extra- and intracellular metabolites. Specific extraction procedures are described below in more detail.

### Methanol and methanol:water extraction

In each case of different methanol extraction (M, MW1, MW2 and MW3), 20 mg of lyophilized pellet powder from each sample was extracted using 1 mL of extraction solution. After sonication for 10 min, the samples were centrifuged at 13000 rpm at 4°C for 5 min and the supernatant was collected. Extraction was repeated by adding another 1 ml of extraction solution to the pellet. After centrifugation, the supernatant from the two extractions were combined, dried by vacuum evaporator and stored at −80°C for further analysis. The extraction procedure was the same for the extracellular metabolites by using rotary shaker for 15 min instead sonication.

### Methanol:chloroform:water extraction

The protocol used for the extraction of the polar and non-polar metabolites from these matrices was adapted from that of Vrhovsek et al. (2012). Briefly, 20 mg of powder from each lyophilized pellet sample was extracted using 1 mL of a mixture of water/methanol/chloroform (20:40:40). After vortexing for 1 min, the samples were put in an orbital shaker for 15 min at room temperature. Samples were centrifuged at 13000 rpm and 4°C for 10 min, and the upper phases constituted of aqueous methanol extract were collected. Extraction was repeated by adding another 600 µL of water/methanol (1:2) to the pellet and chloroform fractions and shaking for 15 min. After centrifugation, the upper phases from the two extractions were combined, dried by vacuum evaporator and stored at −80°C for further analysis. The chloroform phase was also collected in a separated tube, dried by vacuum evaporator and stored at −80°C for the analysis of non-polar metabolites. The same extraction procedure was performed for extracellular metabolites by using rotary shaker for 15 min instead of sonication.

## UPLC-MS analysis and data processing

### Sample Preparation

Dried samples of M, MW1, MW2, MW3 and aqueous fraction of MCW extraction were reconstituted with 400 µl of water (LC-MS grade). Quality control (QC) sample was pooled from 120 µl of each sample and later used for stability and reproducibility assessment of the machine. Pooled QC were diluted to 80%, 60%, 40%, 20%, 10%, 1% and 0% to obtain dilution QC samples. This dilution series were further used in feature extraction step to eliminate product ions with erratic behaviour. Modified from Lewis et al. (2016), the standardized 96-wells plate sample preparation platform were applied in this study. Before aliquoting to the microplate, all samples were randomized in their arrangement in order to reduce technical bias. The 150 µl of samples were then transferred to 96-wells microplate and added up with 150 µl of internal standards (hippuric acid-D5 and L-phenylalanine-13C9, 15N). The mixture was shaken using Thermomixer C (Eppendorf, USA) at 4,500 rpm, 4°C for 10 min. The 125 µl of sample mixture was aliquoted into 2 analytical plates and placed in 4°C sample manager for further analysis in LC/MS positive and negative ionization mode. Whereas, organic phase extracted from methanol/chloroform/water (2:2:1) was resuspended with 400 µl of isopropanol/acetonitrile/water (2:1:1). The 400 µl of lipid internal standards (LPC(9:0), PC(11:0/11:0), FA(17:0), PG(15:0/15:0), PE(15:0/15:0), PS(17:0/17:0), PA(17:0/17:0), Cer(d18:1/17:0), DG(19:0/19:0), PC(23:0/23:0), TG(15:0/15:0/15:0), and TG(17:0/17:0/17:0)) were mixed with 100 µl of samples in 96-wells plate. Subsequent steps of mixing and aliquoting were carried out in accordance with above-mentioned aqueous phase method. The QC and dilution QC samples were also prepared accordingly.

### Metabolite analysis using UPLC-MS

The UPLC-MS running conditions and parameters were slightly modified from (Vorkas et al. 2015). UPLC separation was conducted using an Acquity UPLC system (Waters Corp., USA) connected with HSS T3 (1.8 µm, 2.1×100 mm) column. Column temperature was set at 45°C. Mobile phase A was 0.1% formic acid in water while mobile phase B consisted of 0.1% formic acid in acetonitrile. The elution gradient was set as follows: 99% A (0-0.1 min, 0.4 ml/min), 99-45% A (0.1-10 min, 0.4 ml/min), 45-35% A (10-10.15 min, 0.4-0.41 ml/min), 35-25% A (10.15-10.30 min, 0.41-0.43 ml/min), 25-15% A (10.30-10.45 min, 0.43-0.47 ml/min), 15-5% A (10.45-10.6 min, 0.47-0.55 ml/min), 5-0% A (10.6-10.7 min, 0.55-0.6 ml/min), 0% A (10.7-11 min, 0.6-0.8 ml/min), 0% A (11-12.55 min, 0.8 ml/min), 0-99% A (12.55-12.65 min, 0.8 ml/min), and 99% A (12.65-13.65 min, 0.8-0.4 ml/min). Sample loop of 2 µl was used and injection volume was set at 15 µl. Mass spectrometry was performed using Xevo-G2 XS Q-ToF (Waters Corp., USA) equipped with electrospray ionization (ESI) source. Mass detection was scanned from 50-1200 m/z with scan time at 0.1 s. Source temperature was set at 120°C along with cone gas flow at 150 l/h and desolvation gas flow at 1000 l/h. Cone voltage was 20 V while capillary voltage was 1500 and 1000 V for positive and negative mode, respectively. Leucine enkephalin (200 ng/ml in 50% acetonitrile) was used for lock mass correction with infusion flow rate of 15 µl/min and scan frequency of 60 s. Data was acquired in MS centroid mode using MassLynx software (Waters Corp., USA).

Prior to samples analysis, 5 blanks and 30 pooled QCs were injected to conditioning the column and the instrument and following by 46 dilution QCs among 8 concentrations. Number of injections was 10, 5, 3, 3, 5, 10, and 10 times for 100%, 80%, 60%, 40%, 20%, 10%, 1%, and 0% serial dilutions, respectively. To determine reproducibility of retention time and peak intensity, pooled QC was injected at interval of every 5 samples throughout the entire experiment.

### Lipid profiling using UPLC-MS

UPLC separation was performed using an Acquity UPLC system (Waters Corp., USA) connected with CSH C18 (1.7 µm, 2.1×100 mm) column. Column temperature was set at 55°C. Mobile phase A consisted of water/acetonitrile/isopropanol (2:1:1) plus 20 µM phosphoric acid while mobile phase B was made of isopropanol/acetonitrile (9:1). In both solutions, ammonium acetate was also diluted to 5mM and acetic acid to 0.05%. The elution gradient was set as follows: 99% A (0-0.1 min, 0.4 ml/min), 99-60% A (0.1-2 min, 0.4 ml/min), 60-5% A (2-11.5 min, 0.4 ml/min), 5-0.1% A (11.5-12 min, 0.4-0.45 ml/min), 0.1% A (12-12.5 min, 0.45 ml/min), 0.1-99% A (12.5-12.95 min, 0.45-0.4 ml/min), and 99% A (12.95-14.25 min, 0.4 ml/min). Sample loop of 2 µl was used and injection volume was set at 15 µl to ensure full-loop injection. Mass spectrometry was performed using Xevo-G2 XS Q-ToF (Waters Corp., USA) equipped with electrospray ionization (ESI) source. Mass detection was scanned from 50-2000 m/z with scan time at 0.1 s. Source temperature was set at 120°C along with cone gas flow at 150 l/h and desolvation gas flow at 1000 l/h. Cone voltage was 25 V while capillary voltage was 2000 and 1500 V for positive and negative mode, respectively. Leucine enkephalin (200 ng/ml in 50% acetonitrile) was continuously infused at flow rate of 15 µl/min for mass calibration. Data was collected in centroid mode of normal MS scan using MassLynx software (Waters Corp., USA). Injection order was implemented in the same way as mentioned in metabolite analysis.

### Data processing

Feature extraction, data filtration and quality assessment were undertaken using Progenesis QI software (Nonlinear Dynamics, USA). Raw files of run-order QCs, dilution QCs and all samples were imported into the software and only M+H and M-H adducts were assigned for ion detection in positive and negative mode, respectively. Peak alignment was automatically processed using the most suitable QC as reference and manual adjustments were made where necessary in cases of peak misalignments. Peak picking was also performed by default with minimum peak width of 0.01 min. Data was normalized to internal standards that were able to be detected in acceptable peak shape and intensity. In data filtration step, whole dataset was exported and run using Microsoft Excel 2010 (Microsoft, USA). To yield the greatest extent of final dataset, a relative standard deviation (RSD) of feature intensity among all run-order QC samples and Pearson correlation of features to serial QC dilutions was applied to filter features. Taking all QC samples, the ions with RSD value of ion intensity over 30% were removed from the dataset. As well, the features having a Pearson correlation to dilution series less than 0.8 were considered as artificial features and removed.

### Statistical analysis

All statistical analyses were performed inside the R statistical computing environment (R Core Team 2018) and using the *ggplot2* package for visualisation (Hadley 2015). Principal component analysis (PCA) was performed and visualised on column mean-centered and scaled data using the R packages *FactoMineR* (Lê et al. 2008) and *factoextra* (Kassambara and Mundt 2016). Analysis of variance (ANOVA) were employed on the positive ionization mode data followed by false discovery rate (FDR) correction. The data were log-transformed before the analysis.

## Results and Discussion

### General profile of metabolites in two lab-scale EBPR bioreactors

We first examined the differences in metabolite profiles among samples from each reactor using PCA categorised by extraction method and additionally by compartment (intracellular or extracellular) in the case of Reactor B (Fig. 1). In Reactor A, we observed that different extraction methods induced distinct metabolite profiles were in both positive (Fig 1A) and negative ionization mode (Fig 1C), and that samples from the same extraction methods and physiological state were clustered together. Plotting PCA sample scores confirmed that MW1 and MW2 presented related profiles in negative mode and the best separation of metabolites in anaerobic and aerobic stages. For Reactor B, three extraction methods for extracellular metabolites were less distinct from each other in positive mode (Fig. 1B) but better differentiated in negative mode while intracellular metabolites were visibly clustered both in positive and negative mode. Moreover, the profiles of extracellular metabolites are clearly distinguished from intracellular metabolites (Fig. 1D).

The data from non-polar phase derived from the MCW extraction method were also investigated for sample to sample variation of each time point in Reactor A and each sample type in Reactor B. According to the PCA score plot (Supplementary Material 2 - Fig. 3), the MCW could not give a clear separation among the different time points in Reactor A. However, MCW extraction for Reactor B apparently showed the large variation between extra- and intra-cellular non-polar metabolites with overlapping data points between two different stages of time during the EBPR cycle (Supplementary Material 2 - Fig 3).

### Selection of optimal extraction method

To identify the optimal solvent system for the global metabolite profiling of microbial communities in reactors, we evaluated the performance of five extraction methods based on three criteria; *1*) the uniformity of mass features distribution across the retention time axis, *2*) the peak intensities of metabolites and *3*) the number of differentially expressed mass features between the time points during the EBPR cycle and sample types, under the hypothesis that many metabolite profiles should show differential abundance levels given the transition between anoxic and aerobic physiological states across the course of the cycle study.

In Reactor A, MW1 generated the most uniform distribution of peaks across the retention axis, followed by MW2, whereas M gave the lowest diversity of peaks. Total ion chromatogram of metabolite profiles from each extraction method in Reactor A are shown in Supplementary Material 4. The second factor we considered was the peak intensity of each metabolite. The map of feature positions highlighted with highest mean values of each extraction method can be seen in Supplementary Material 2 - Fig 4. MW1 methods generated a greater number of peaks, and with higher signal level, than the other four methods. The third criterion for selection of optimal extraction solvent is the number of metabolite features statistically significantly different from the different experimental condition. Finally, as shown in Supplementary Material 3 - Fig 5, ANOVA analysis highlighted that MW1 gave the largest number of differentially abundant mass features among different stages during the EBPR cycle. In regards the MCW method, the number of peaks detected in the aqueous phase were less than observed with methanol:water extraction methods, and this may be due to partitioning of some non-polar metabolites into the chloroform phase. However, the peak intensities were the lowest among the extraction methods and no significant differences among time points were observed by MCW. These data indicate that MW1 is the most appropriate solvent for the extraction of metabolites from biomass in Reactor A.

Unlike the case of Reactor A, MW1 was not suitable for extracting the metabolome of microbial communities resident in Reactor B. The metabolite profiles indicated the MCW method contained the largest number of peaks in total ion chromatograms among the different extraction methods (Supplementary Material 4 and Table 1), yielding the best uniformity of mass feature distribution over retention time with the largest number of peaks Supplementary Material 3 - Fig 6. In addition, the largest number of differentially expressed mass features between anaerobic and aerobic stages by MCW extraction method was confirmed by ANOVA analysis (Supplementary Material 3 - Fig 7). From the chloroform phase data, the high number of peaks (Table 1) and increased number of differentially abundant features (Supplementary Material 3 - Fig 7), it is likely that the microbial community from Reactor B is lipid-rich, which may include a complex mixture of soluble microbial products (SMPs) (Barker and Stuckey 1999). The MCW method would be better suited to this type of community biomass, given the potential for better separation of polar and non-polar metabolites. In a recent study, Tipthara and colleagues applied the extraction method in an analysis of metabolite and lipid SMPs from the supernatant of an anaerobic digester batch reactor, but using different volume ratios of extraction solvents (1:2:1) (Tipthara et al. 2017).

**Table 1.** Comparison of total detected peak numbers extracted from different solvent extraction methods (positive and negative mode) for Reactor A and Reactor B.

| Extraction Method | Total number of detected peaks |  |  |  |
| --- | --- | --- | --- | --- |
|  | Reactor A |  | Reactor B |  |
|  | Positive mode | Negative mode | Positive mode | Negative mode |
| M<sup>a</sup> | 6934 | 2804 | 1262 | 1428 |
| MW1<sup>b</sup> | 8183 | 4045 | 1334 | 1816 |
| MW2<sup>c</sup> | 7699 | 3852 | N/A | N/A |
| MW3<sup>d</sup> | 7941 | 3604 | N/A | N/A |
| MCW-aqueous fraction<sup>e</sup> | 7825 | 3702 | 1364 | 1735 |
| MCW -<br>chloroform fraction<sup>e</sup> | 2595 | 527 | 3507 | 514 |

<sup>a</sup>M: pure methanol
<sup>b</sup>MW1: methanol:water (50:50 v/v)
<sup>c</sup>MW2: methanol:water (60:40 v/v)
<sup>d</sup>MW3: methanol:water (80:20 v/v)
<sup>e</sup>MCW: methanol:chloroform:water (40:40:20 v/v/v)

In untargeted microbial metabolomics studies, the chosen extraction method is determined mostly by the physicochemical properties of a given biological sample, such as polarity, selectivity, toxicity and inertness (Kim and Verpoorte 2010). Cold methanol extraction solvent is used to simultaneously quench and extract metabolites, having the advantage of being a rapid process and permitting the recovery of a large variety of metabolites. It has been widely used for extraction of intracellular metabolites from animal (Dettmer et al. 2011), bacterial (Maharjan and Ferenci 2003) and fungal cells (Villas-Bôas et al. 2005). For example, when methanol:water mixture was applied for extraction of intracellular metabolites in *Escherichia coli*, (Kimball and Rabinowitz 2006) found that the volume ratio of methanol and water solvent impacted the effectiveness of extraction and recommended 80:20 v/v methanol:water. In contrast, a 50:50 v/v ratio was also chosen for *Nanoarcheum equitans* and its archaeal host *Ignicoccus hospitalis* (Hamerly et al. 2015) and for *Pseudomonas fluorescens, Streptomyces coelicolor* and *Saccharomyces cerevisiae* (Villas-Bôas and Bruheim 2007). Such species specificity implies challenges for undertaking extraction procedures in the context of microbial communities and microbiomes, which includes the presence of extra polymeric substance (EPS) and spatially clustered biomass, such as floccular or granular morphology. This is consistent with the findings of the present study and implies that extraction method appears highly biomass-specific and needs to be optimized accordingly.

## Conclusion

We used untargeted metabolomics analysis to investigate optimal metabolite extraction solvents for different microbial communities in activated sludge enrichment reactors, demonstrating that extraction methods need to be carefully tailored based on the microbial community under study, even among communities with similar phenotypes. Our approach provides direct surveys of the metabolic state of PAO-enriched EBPR communities, and these data build the foundation for ongoing integrative omics studies of these complex microbial communities.

## Supporting information

Supplementary Data for Reactor A

Supplementary Data for Reactor B

## Acknowledgements

This research was supported by the Singapore National Research Foundation and the Ministry of Education under the Research Centre of Excellence Programme, and by a research grant from the National Research Foundation under the Environment and Water Industry Programme (Project number: 1102–IRIS–10–02). We thank Dr Sean Ng and Professor Jeremy Everett for their contributions to the early stages of this work.

## Data Availability

All raw data from this study has been deposited at the MetaboLights data respository (accesion numbers pending at the time of writing).

## Author Contributions

NMMTS and RBHW conceived the study and designed analyses. GQ, REZM and SW established and maintained enrichment reactor communities. NMMTS, GQ and REZM performed reactor experiments and obtained samples. NMMTS performed extractions. PS carried out the UPLC-MS assays and performed raw data processing. NMMTS performed all subsequent data analysis. The manuscript was mainly written by NMMTS with contributions from all other authors.

## Compliance with ethical standards

### Conflict of interest

The authors declare that they have no conflict of interest.

### Ethical approval

This article does not contain any studies with human participants or animals performed by any of the authors.

## List of Supplementary Material

Supplementary Method 1 - Operation of lab-scale enrichment bioreactors

Supplementary Material 2 - Figures

Supplementary Material 3 - TIC plots_reactor A

Supplementary Material 4 - TIC plots_reactor B

