## Supplementary Methods for "Evaluation of extraction solvents for untargeted metabolomics analysis of enrichment reactor cultures performing enhanced biological phosphorus removal (EBPR)"

**Operation of the lab-scale enrichment bioreactors (Saw *et al.*)**

**Reactor A**

Reactor A was a sequencing batch reactor (SBR) with 5.4 L working volume, and was inoculated with activated sludge obtained from an existing EBPR enrichment reactor, with a protocol designed to enrich for known PAO species from the genus *Candidatus* *Accumulibacter*. The SBR was operated with 6 h cycles, including a feeding (60 min) stage, an anaerobic stage (20 min), an aerobic stage (180 min), and a settling/decant stage (100 min). In each cycle, 2.35 L of synthetic wastewater composed of 0.53 L of solution A (containing 1.02g/L NH_4_Cl, 1.2g/L MgSO_4_ 7H_2_O, 0.01g/L peptone, 0.01g/L yeast extract and 6.8 g/L sodium acetate) and 1.82L of solution B (0. 312g/L K_2_HPO_4_ 3H_2_O, 0.185 g/L KH_2_PO_4_, 0.75 mg/L FeCl_3_ 6H_2_O 0.015 mg/L CuSO_4_ 5H_2_O, 0.03 mg/L MnCl_2_, 0.06 mg/L ZnSO_4_, 0.075 mg/L CoCl_2_, 0.075 mg/L H_3_BO_3_, 0.09mg/L KI and 0.06 mg/L Na_2_MoO_4_ 2H_2_O; modified from Lu et al. 2006) were introduced into the reactor continuously (in 60 min). The reactor was operated at 31^o^C with a HRT and a SRT of 12 h and 7 days, respectively. The pH was controlled at 7.20-7.60 with DO levels maintained at 0.8-1.2 mg/L during the aerobic phase.

**Reactor B**

Reactor B was a 5-L SBR containing an EBPR microbial consortia was fed with synthetic wastewater containing glucose as the main carbon source. The cycle duration was 6 h and consisted of 30 min feeding, 125 min anaerobic, 154 min aerobic, 1 min sludge discharge, 35 min settling and 15 min supernatant discharge phases. The HRT was 15 h. The feed composition was adapted from that of Lu et al. (2006) and split into 0.3 and 1.7 L of solutions A and B, respectively. Solution A contained (per litre): 1,852.8 mg glucose anhydrous, 637.5 mg NH4Cl, 6.25 mg peptone, 6.25 mg yeast extract, 750 mg MgSO4·7H2O, 118.75 mg CaCl2·2H2O and 401.6 mg N-Allylthiourea to inhibit nitrification. Feed solution B contained (per litre): 73.20 mg K2HPO4·3H2O, 43.65 mg KH2PO4, 0.55 mL trace elements solution 1 and 0.55 mL trace elements solution 2. The trace elements solutions included (per litre and adapted from Smolders et al. 1994): solution 1: 1.5 g FeCl3·6H2O, 0.03 g CuSO4·5H2O, 0.12 g MnCl2·4H2O, 0.12 g ZnSO4·7H2O, 0.15 g CoCl2·6H2O and 0.1 g EDTA disodium salt, and solution 2: 0.15 g H3BO3, 0.18 g KI and 0.06 g Na2MoO4·2H2O. Nitrogen gas was sparged into the reactor during the anaerobic phase, while air was supplied during the aerobic phase to maintain a dissolved oxygen concentration between 0.5 and 1 mg/L. The temperature was set at 31±1 C and pH was controlled at 7.5±0.25 by using 0.5 M NaOH and 0.5 M HCl.
