## Supplementary Data for Reactor A for "Evaluation of extraction solvents for untargeted metabolomics analysis of enrichment reactor cultures performing enhanced biological phosphorus removal (EBPR)"

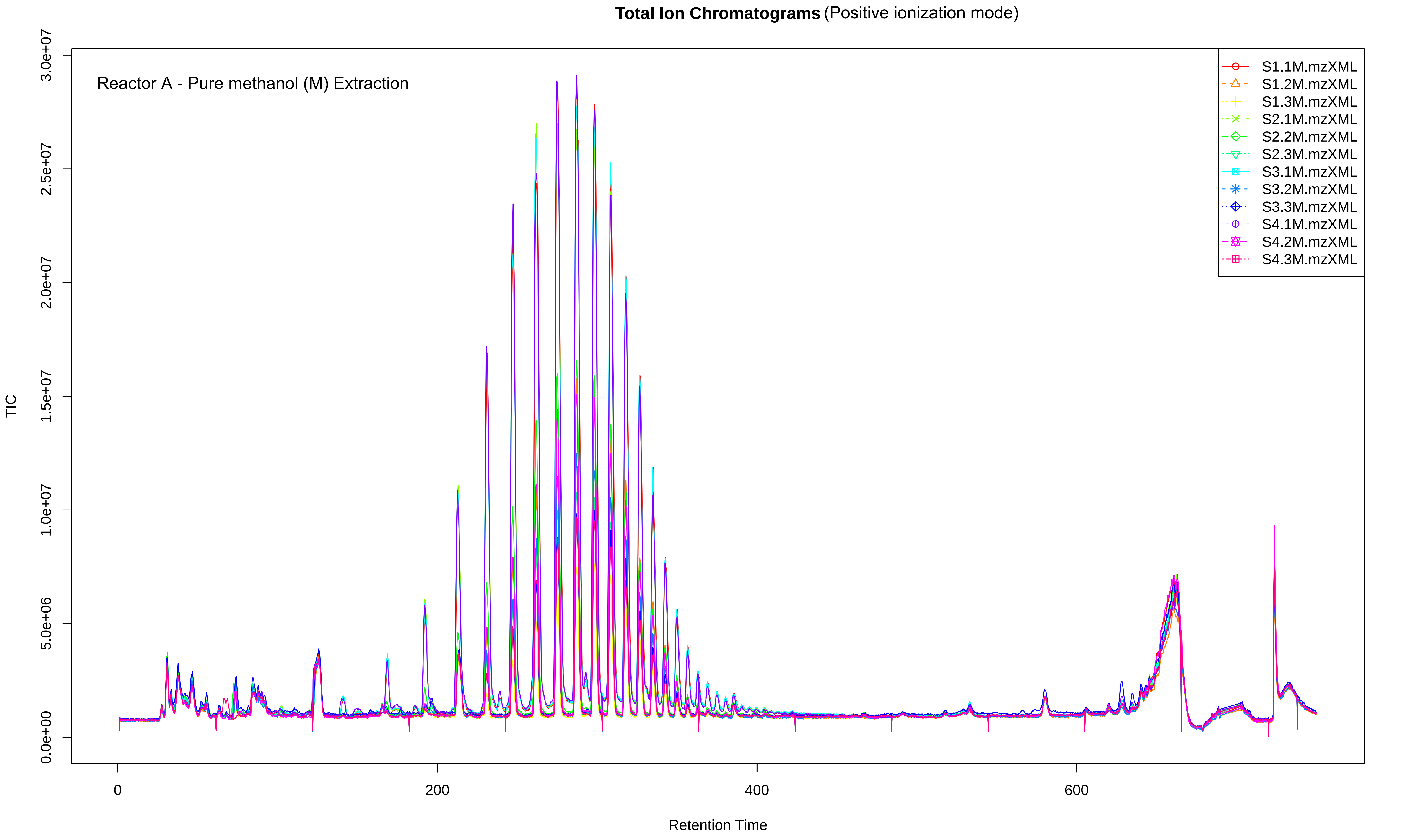
Total Ion Chromatograms (Positive ionization mode)
Reactor A - Pure methanol (M) Extraction
S1.1M.mzXML
S1.2M.mzXML
S1.3M.mzXML
S2.1M.mzXML
S2.2M.mzXML
S2.3M.mzXML
S3.1M.mzXML
S3.2M.mzXML
S3.3M.mzXML
S4.1M.mzXML
S4.2M.mzXML
S4.3M.mzXML
TIC
0.0e+00
5.0e+06
1.0e+07
1.5e+07
2.0e+07
2.5e+07
3.0e+07
0
200
400
600
Retention Time

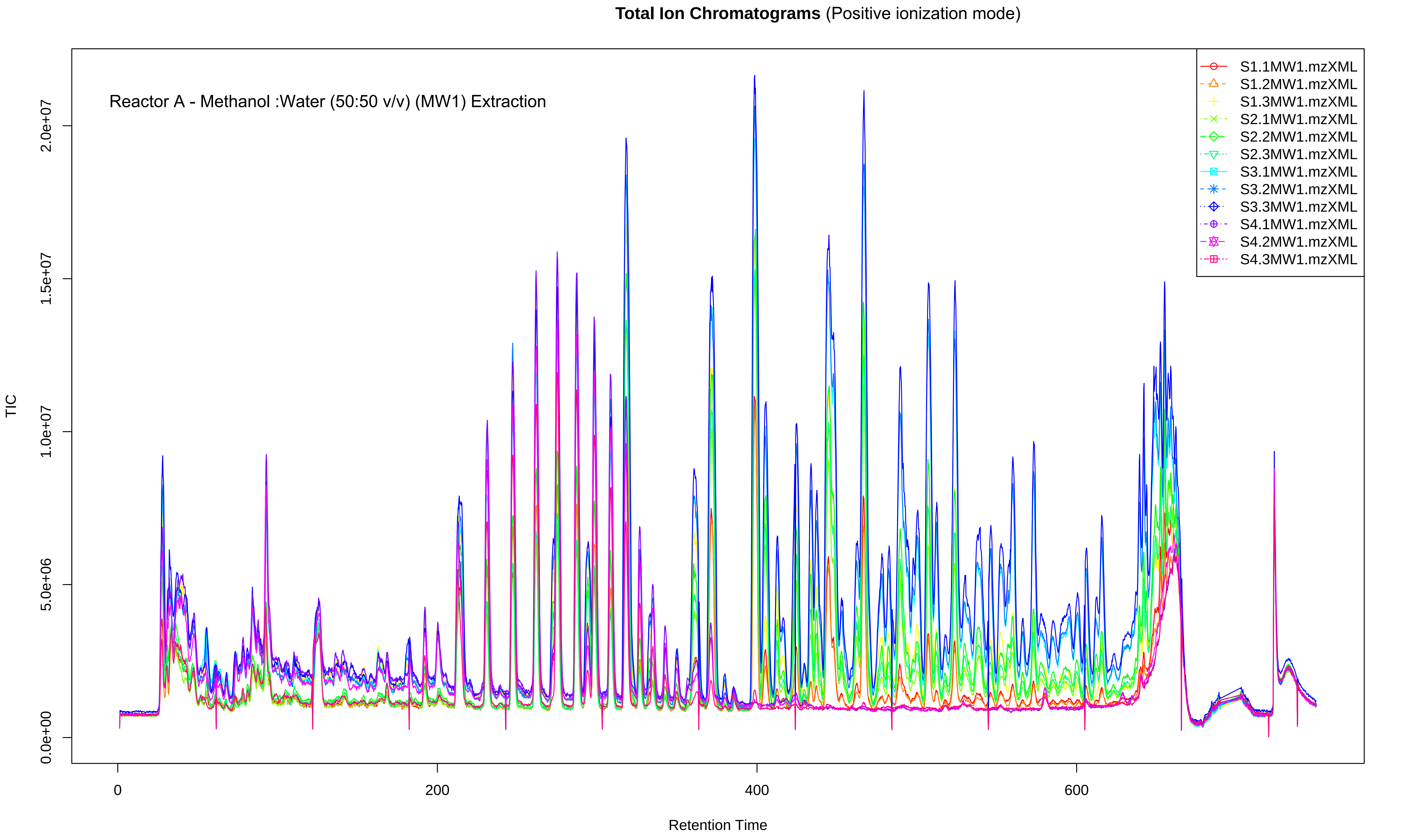

**Total Ion Chromatograms** (Positive ionization mode)
Reactor A - Methanol :Water (50:50 v/v) (MW1) Extraction
TIC
Retention Time
0.0e+00
5.0e+06
1.0e+07
1.5e+07
2.0e+07
0
200
400
600
S1.1MW1.mzXML
S1.2MW1.mzXML
S1.3MW1.mzXML
S2.1MW1.mzXML
S2.2MW1.mzXML
S2.3MW1.mzXML
S3.1MW1.mzXML
S3.2MW1.mzXML
S3.3MW1.mzXML
S4.1MW1.mzXML
S4.2MW1.mzXML
S4.3MW1.mzXML

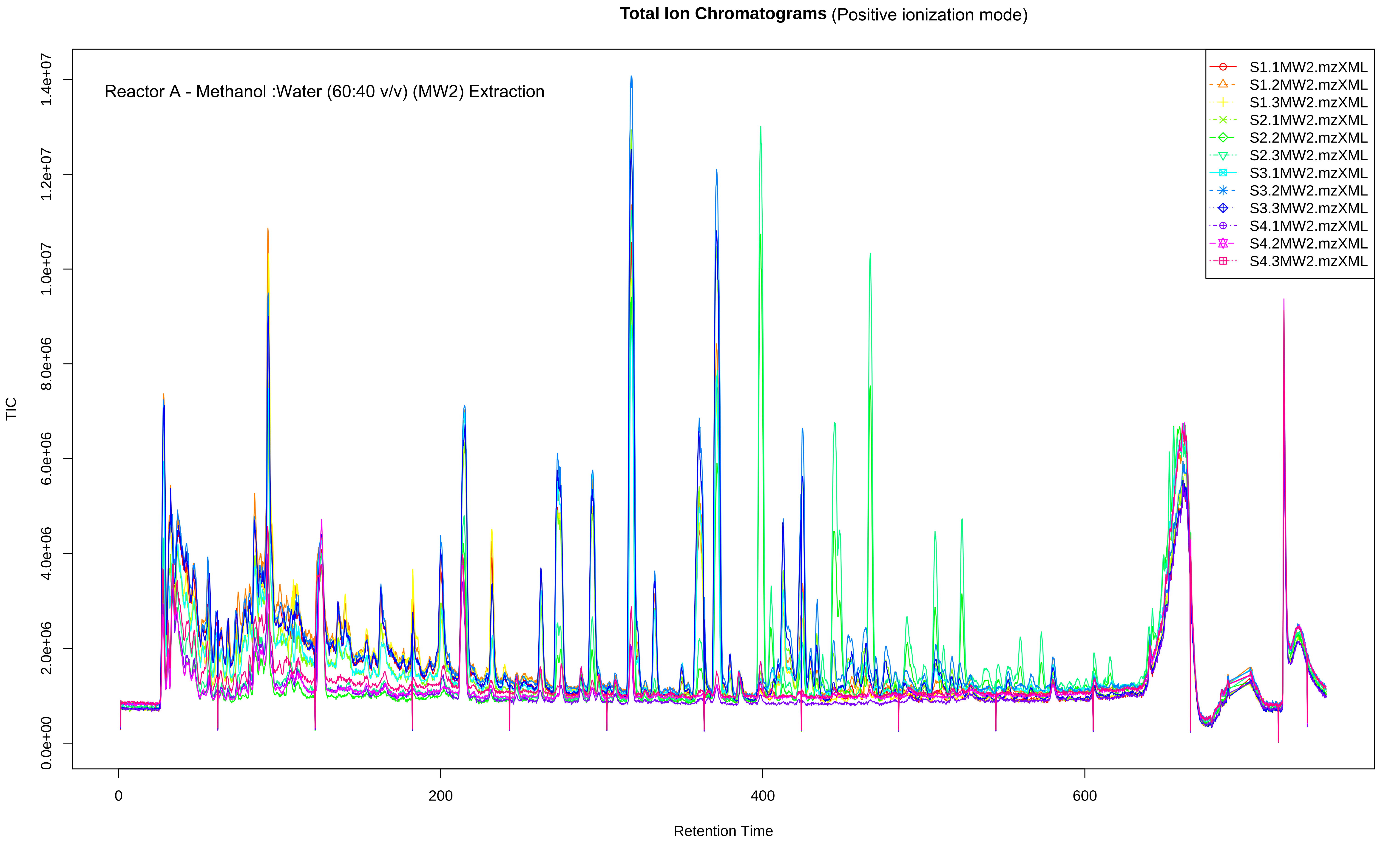
Total Ion Chromatograms (Positive ionization mode)
Reactor A - Methanol :Water (60:40 v/v) (MW2) Extraction
S1.1MW2.mzXML
S1.2MW2.mzXML
S1.3MW2.mzXML
S2.1MW2.mzXML
S2.2MW2.mzXML
S2.3MW2.mzXML
S3.1MW2.mzXML
S3.2MW2.mzXML
S3.3MW2.mzXML
S4.1MW2.mzXML
S4.2MW2.mzXML
S4.3MW2.mzXML
TIC
0.0e+00
2.0e+06
4.0e+06
6.0e+06
8.0e+06
1.0e+07
1.2e+07
1.4e+07
0
200
400
600
Retention Time

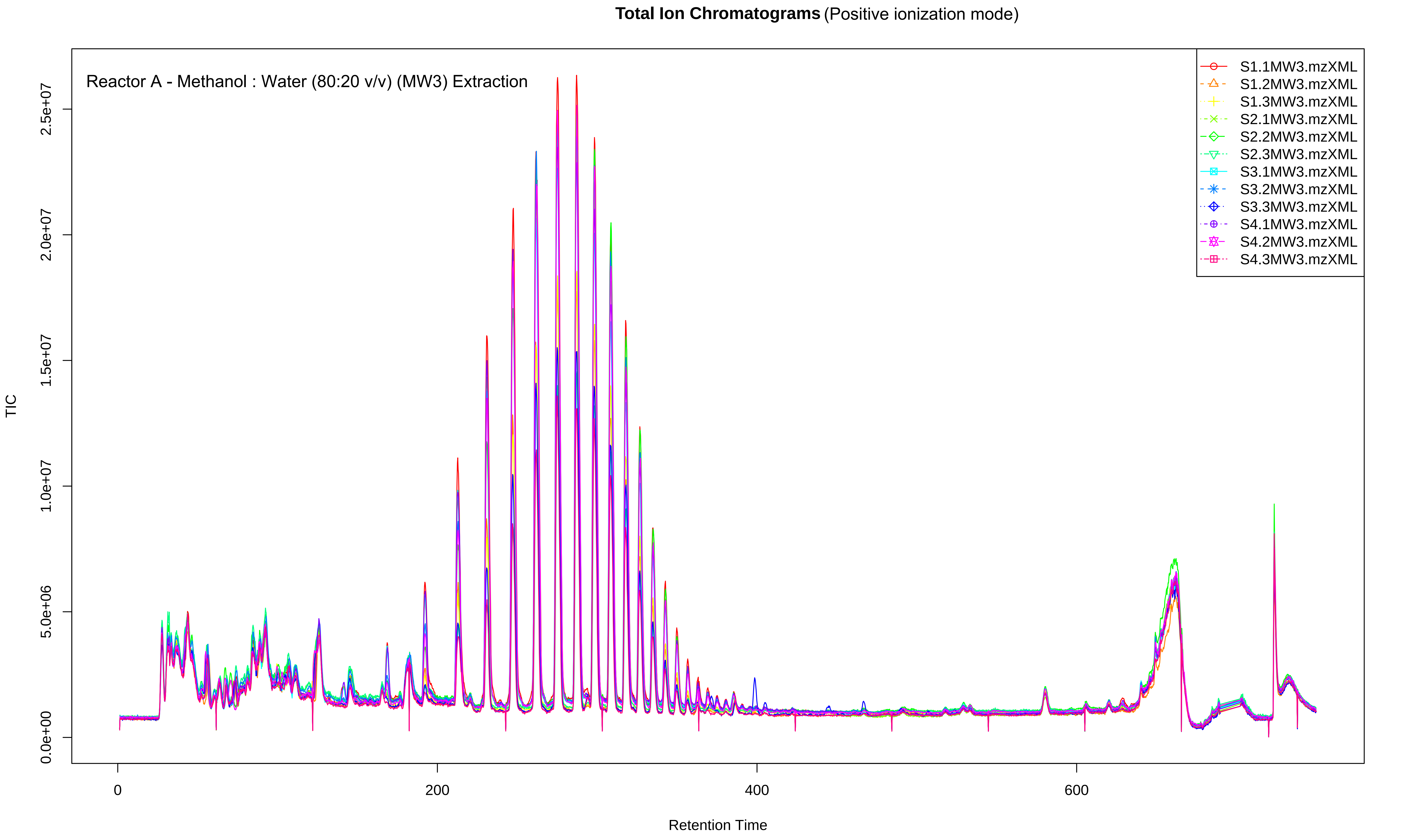

Total Ion Chromatograms (Positive ionization mode)
Reactor A - Methanol : Water (80:20 v/v) (MW3) Extraction
TIC
Retention Time
0.0e+00
5.0e+06
1.0e+07
1.5e+07
2.0e+07
2.5e+07
0
200
400
600
S1.1MW3.mzXML
S1.2MW3.mzXML
S1.3MW3.mzXML
S2.1MW3.mzXML
S2.2MW3.mzXML
S2.3MW3.mzXML
S3.1MW3.mzXML
S3.2MW3.mzXML
S3.3MW3.mzXML
S4.1MW3.mzXML
S4.2MW3.mzXML
S4.3MW3.mzXML

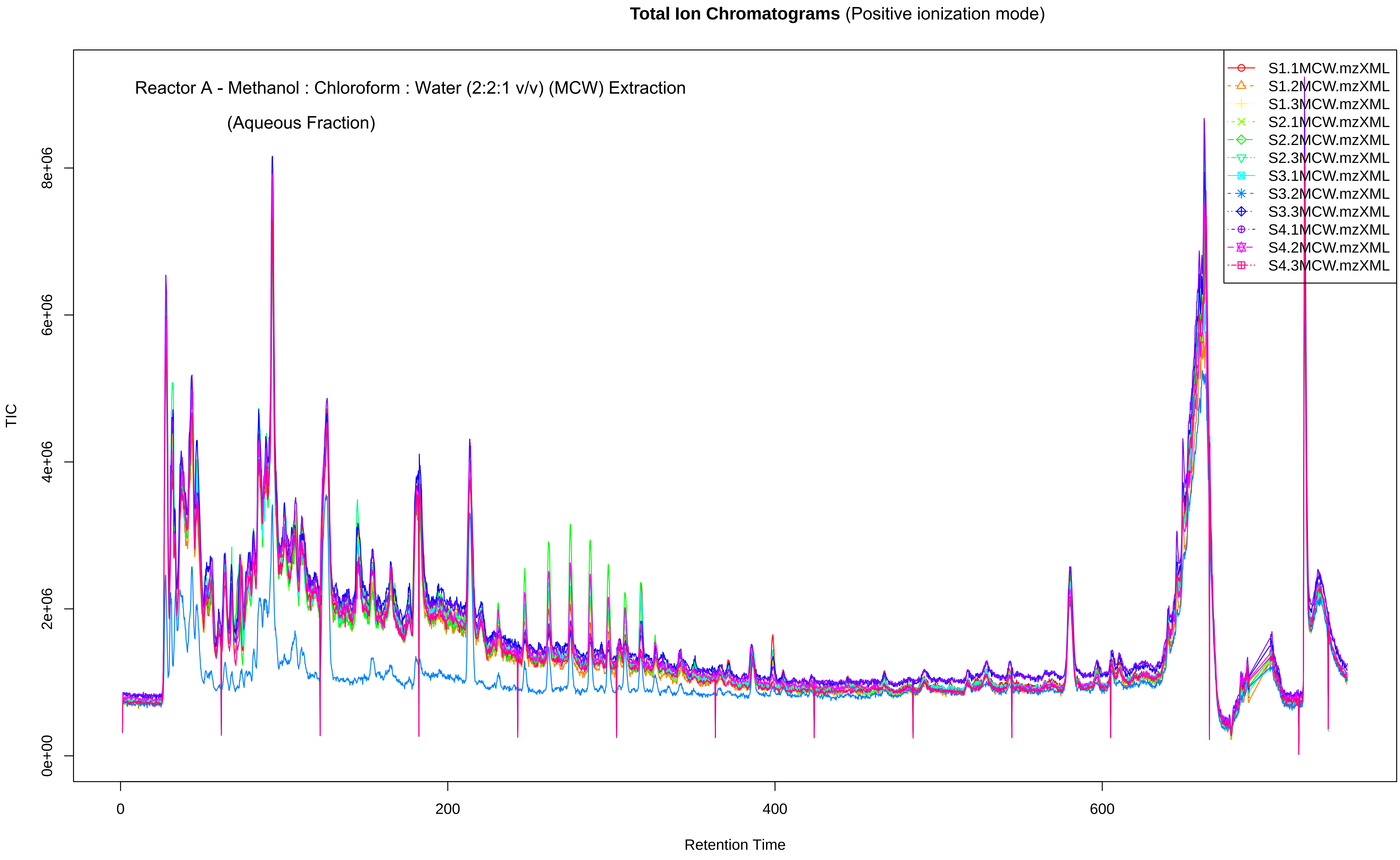

Total Ion Chromatograms (Positive ionization mode)
Reactor A - Methanol : Chloroform : Water (2:2:1 v/v) (MCW) Extraction
(Aqueous Fraction)
S1.1MCW.mzXML
S1.2MCW.mzXML
S1.3MCW.mzXML
S2.1MCW.mzXML
S2.2MCW.mzXML
S2.3MCW.mzXML
S3.1MCW.mzXML
S3.2MCW.mzXML
S3.3MCW.mzXML
S4.1MCW.mzXML
S4.2MCW.mzXML
S4.3MCW.mzXML
TIC
0e+00
2e+06
4e+06
6e+06
8e+06
0
200
400
600
Retention Time

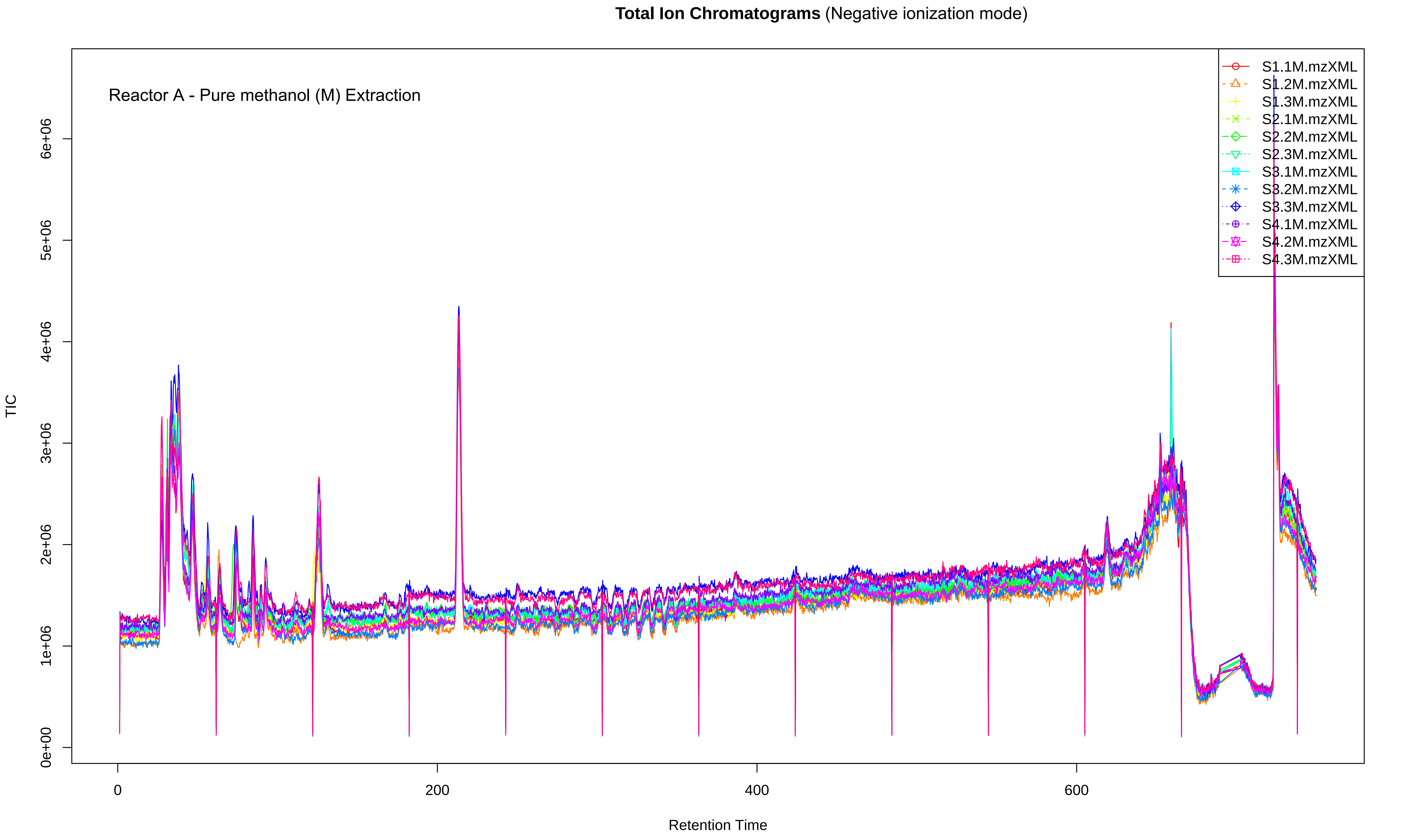

Total Ion Chromatograms (Negative ionization mode)
Reactor A - Pure methanol (M) Extraction
TIC
Retention Time
0e+00
1e+06
2e+06
3e+06
4e+06
5e+06
6e+06
0
200
400
600
S1.1M.mzXML
S1.2M.mzXML
S1.3M.mzXML
S2.1M.mzXML
S2.2M.mzXML
S2.3M.mzXML
S3.1M.mzXML
S3.2M.mzXML
S3.3M.mzXML
S4.1M.mzXML
S4.2M.mzXML
S4.3M.mzXML

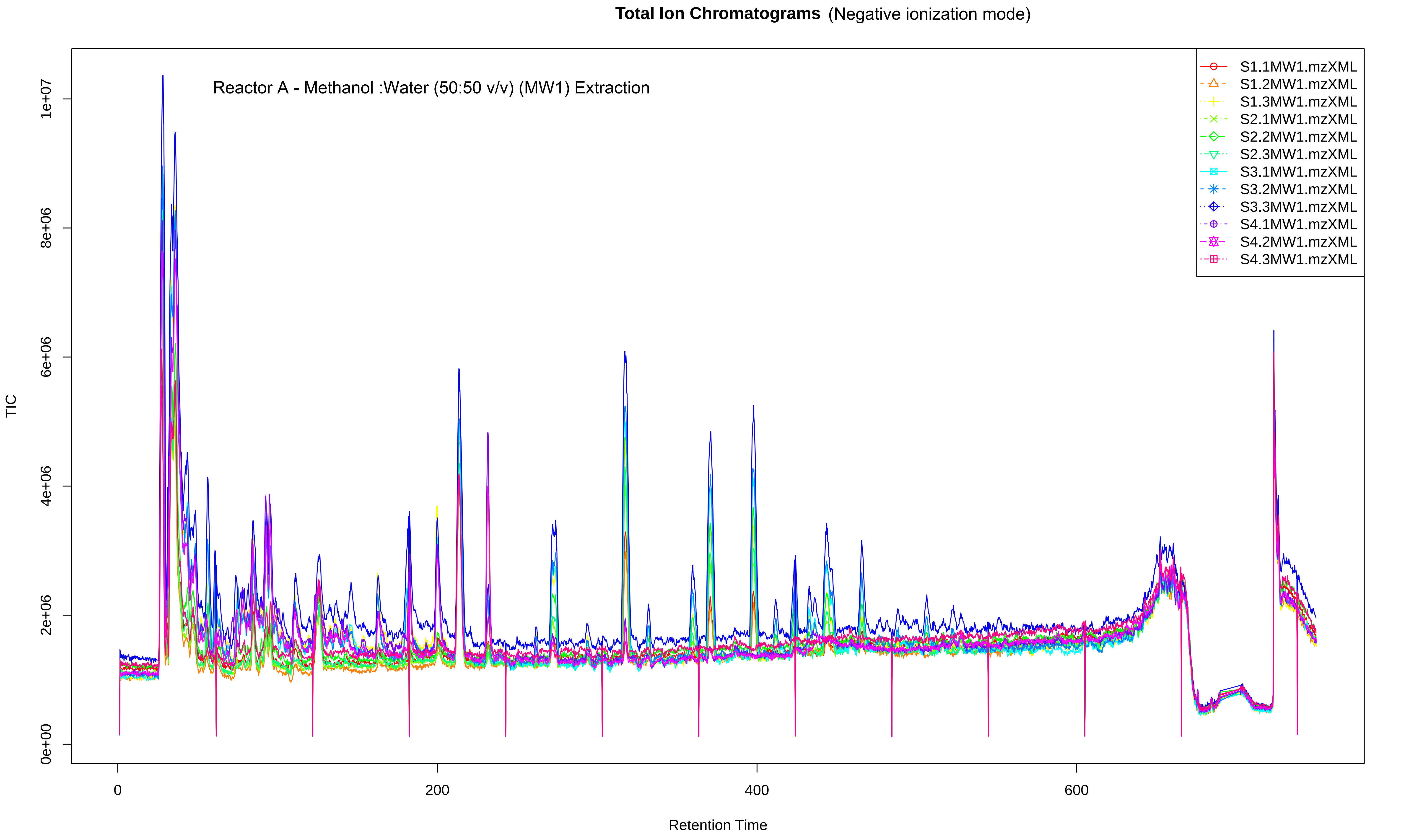

Total Ion Chromatograms (Negative ionization mode)
Reactor A - Methanol :Water (50:50 v/v) (MW1) Extraction
S1.1MW1.mzXML
S1.2MW1.mzXML
S1.3MW1.mzXML
S2.1MW1.mzXML
S2.2MW1.mzXML
S2.3MW1.mzXML
S3.1MW1.mzXML
S3.2MW1.mzXML
S3.3MW1.mzXML
S4.1MW1.mzXML
S4.2MW1.mzXML
S4.3MW1.mzXML
TIC
0e+00
2e+06
4e+06
6e+06
8e+06
1e+07
0
200
400
600
Retention Time

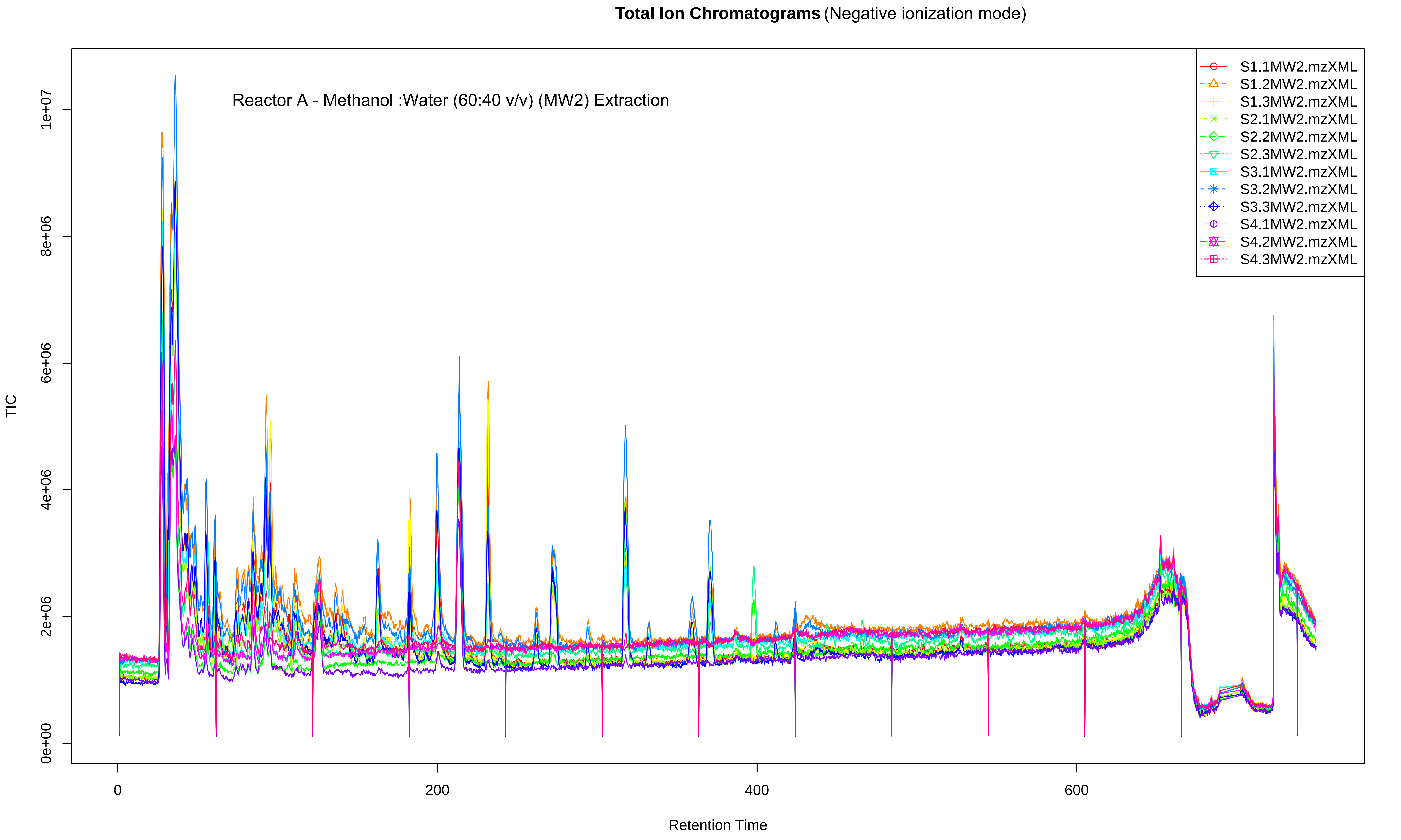

Total Ion Chromatograms (Negative ionization mode)
Reactor A - Methanol :Water (60:40 v/v) (MW2) Extraction
TIC
Retention Time
0e+00
2e+06
4e+06
6e+06
8e+06
1e+07
0
200
400
600
S1.1MW2.mzXML
S1.2MW2.mzXML
S1.3MW2.mzXML
S2.1MW2.mzXML
S2.2MW2.mzXML
S2.3MW2.mzXML
S3.1MW2.mzXML
S3.2MW2.mzXML
S3.3MW2.mzXML
S4.1MW2.mzXML
S4.2MW2.mzXML
S4.3MW2.mzXML

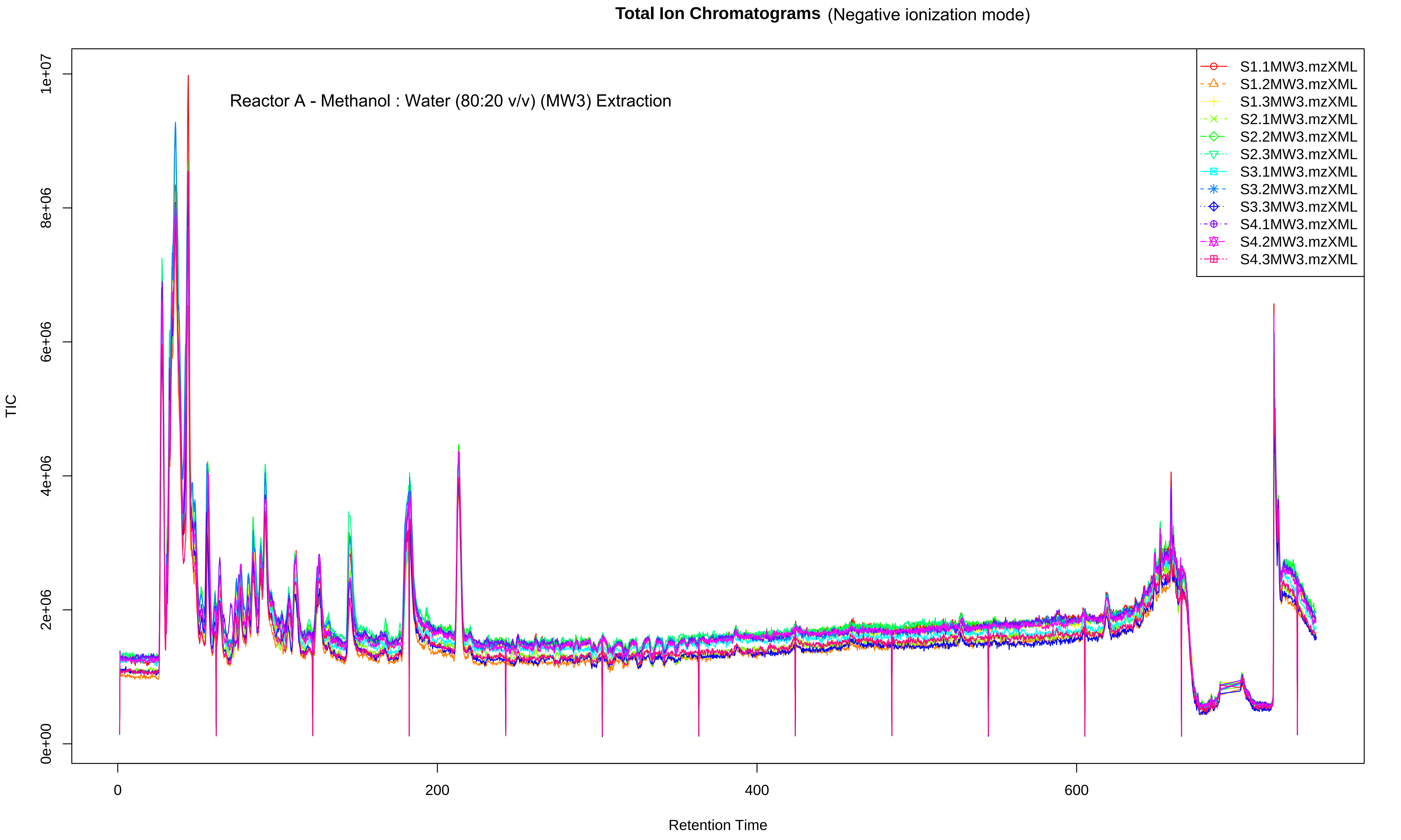
Total Ion Chromatograms (Negative ionization mode)
Reactor A - Methanol : Water (80:20 v/v) (MW3) Extraction
S1.1MW3.mzXML
S1.2MW3.mzXML
S1.3MW3.mzXML
S2.1MW3.mzXML
S2.2MW3.mzXML
S2.3MW3.mzXML
S3.1MW3.mzXML
S3.2MW3.mzXML
S3.3MW3.mzXML
S4.1MW3.mzXML
S4.2MW3.mzXML
S4.3MW3.mzXML
TIC
0e+00
2e+06
4e+06
6e+06
8e+06
1e+07
0
200
400
600
Retention Time

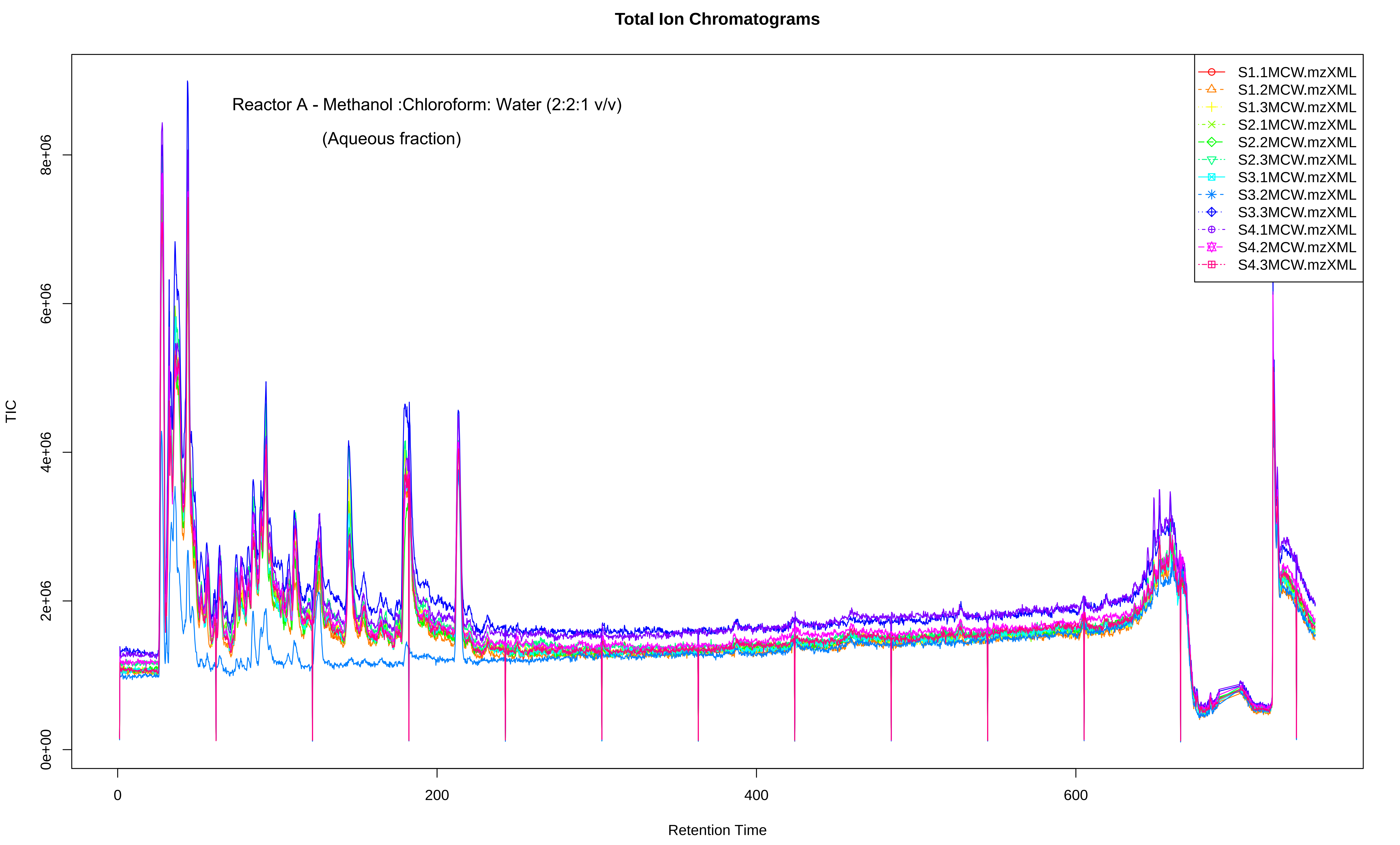
Total Ion Chromatograms
Reactor A - Methanol :Chloroform: Water (2:2:1 v/v)
(Aqueous fraction)
S1.1MCW.mzXML
S1.2MCW.mzXML
S1.3MCW.mzXML
S2.1MCW.mzXML
S2.2MCW.mzXML
S2.3MCW.mzXML
S3.1MCW.mzXML
S3.2MCW.mzXML
S3.3MCW.mzXML
S4.1MCW.mzXML
S4.2MCW.mzXML
S4.3MCW.mzXML
TIC
0e+00
2e+06
4e+06
6e+06
8e+06
0
200
400
600
Retention Time

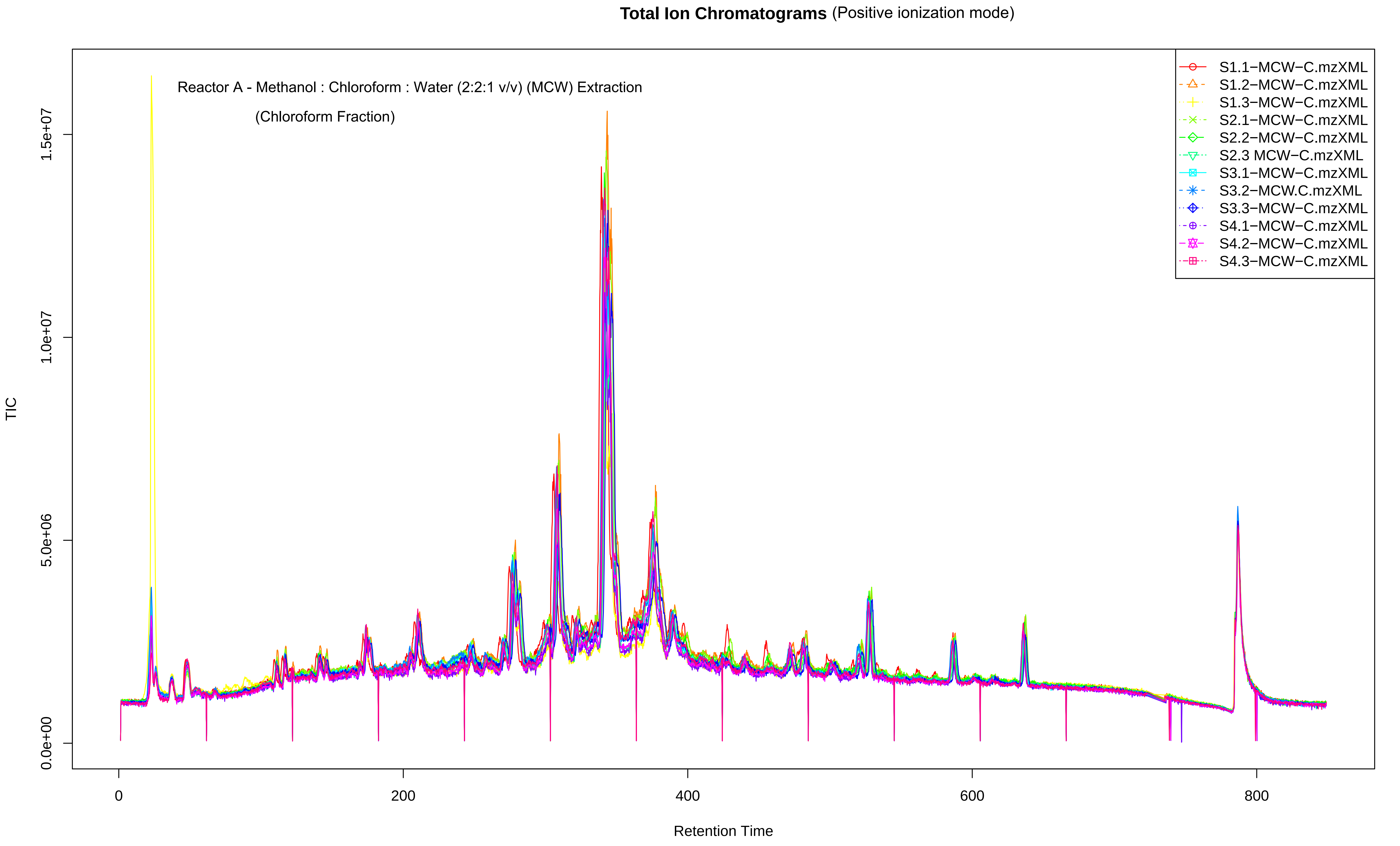
Total Ion Chromatograms (Positive ionization mode)
Reactor A - Methanol : Chloroform : Water (2:2:1 v/v) (MCW) Extraction
(Chloroform Fraction)
TIC
Retention Time
0.0e+00
5.0e+06
1.0e+07
1.5e+07
0
200
400
600
800
S1.1−MCW−C.mzXML
S1.2−MCW−C.mzXML
S1.3−MCW−C.mzXML
S2.1−MCW−C.mzXML
S2.2−MCW−C.mzXML
S2.3 MCW−C.mzXML
S3.1−MCW−C.mzXML
S3.2−MCW.C.mzXML
S3.3−MCW−C.mzXML
S4.1−MCW−C.mzXML
S4.2−MCW−C.mzXML
S4.3−MCW−C.mzXML

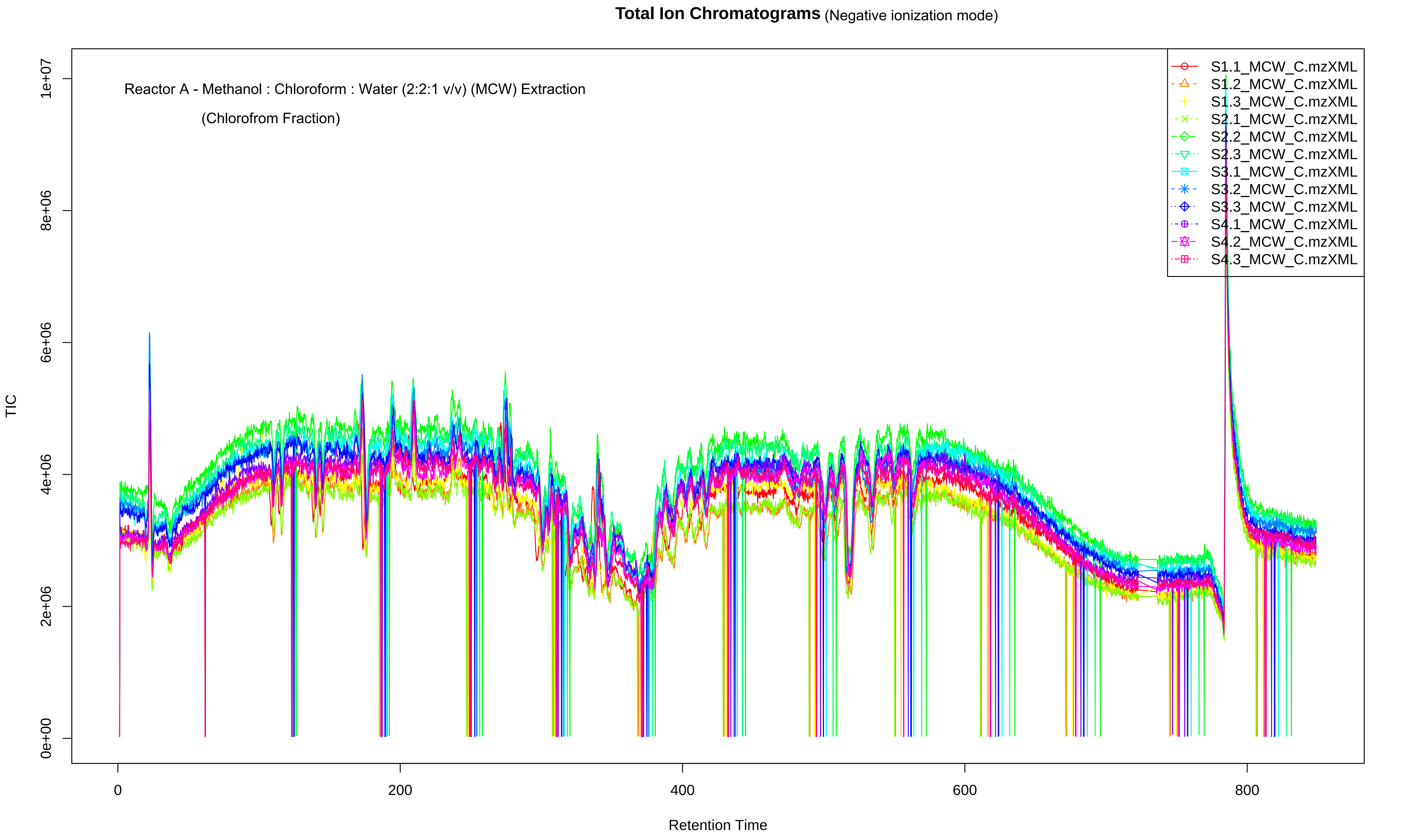

Total Ion Chromatograms (Negative ionization mode)
Reactor A - Methanol : Chloroform : Water (2:2:1 v/v) (MCW) Extraction
(Chlorofrom Fraction)
S1.1_MCW_C.mzXML
S1.2_MCW_C.mzXML
S1.3_MCW_C.mzXML
S2.1_MCW_C.mzXML
S2.2_MCW_C.mzXML
S2.3_MCW_C.mzXML
S3.1_MCW_C.mzXML
S3.2_MCW_C.mzXML
S3.3_MCW_C.mzXML
S4.1_MCW_C.mzXML
S4.2_MCW_C.mzXML
S4.3_MCW_C.mzXML
TIC
0e+00
2e+06
4e+06
6e+06
8e+06
1e+07
0
200
400
600
800
Retention Time
