## Supplementary figures and images for "Evaluation of extraction solvents for untargeted metabolomics analysis of enrichment reactor cultures performing enhanced biological phosphorus removal (EBPR)"

### Supplementary Data for Reactor B

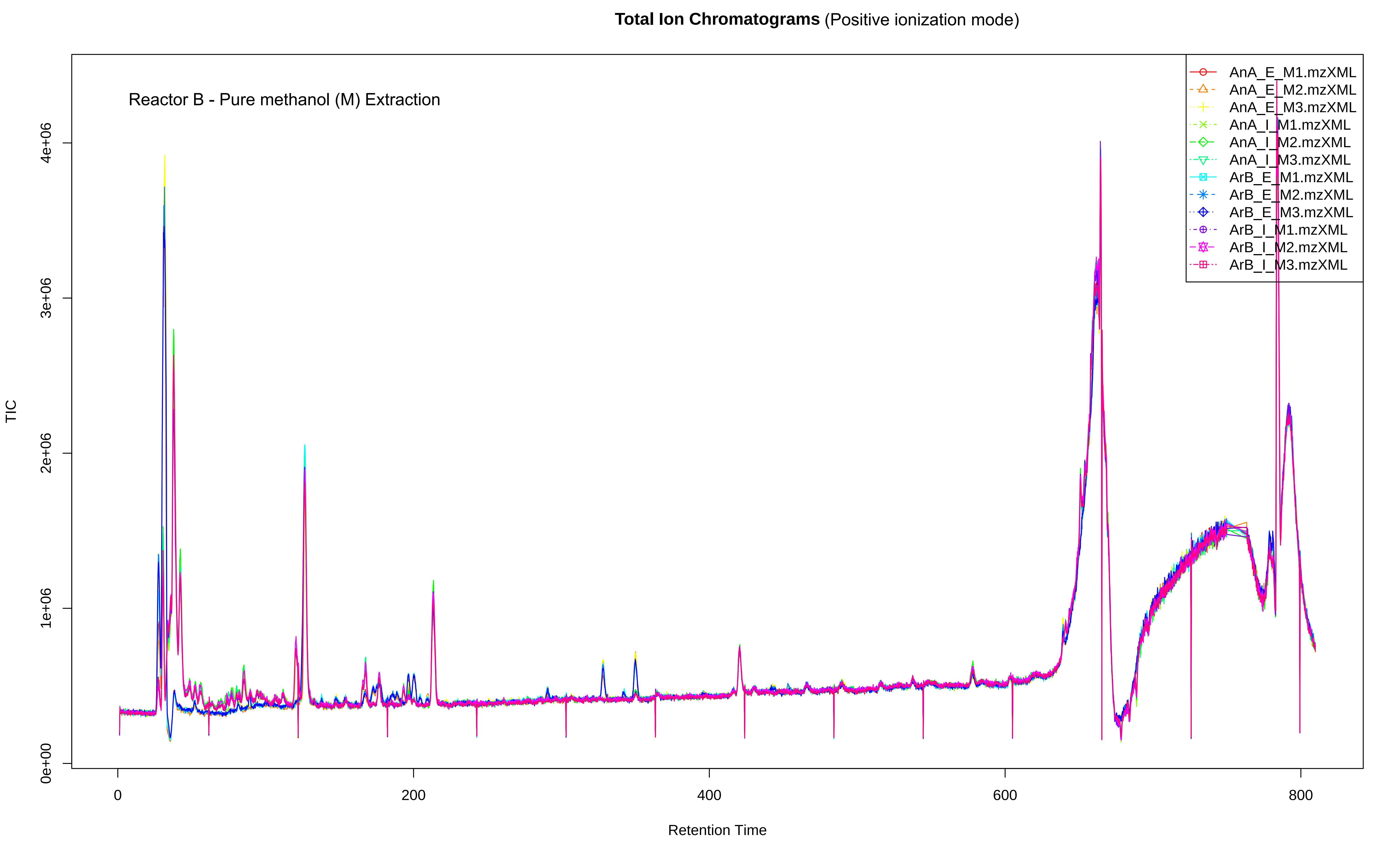

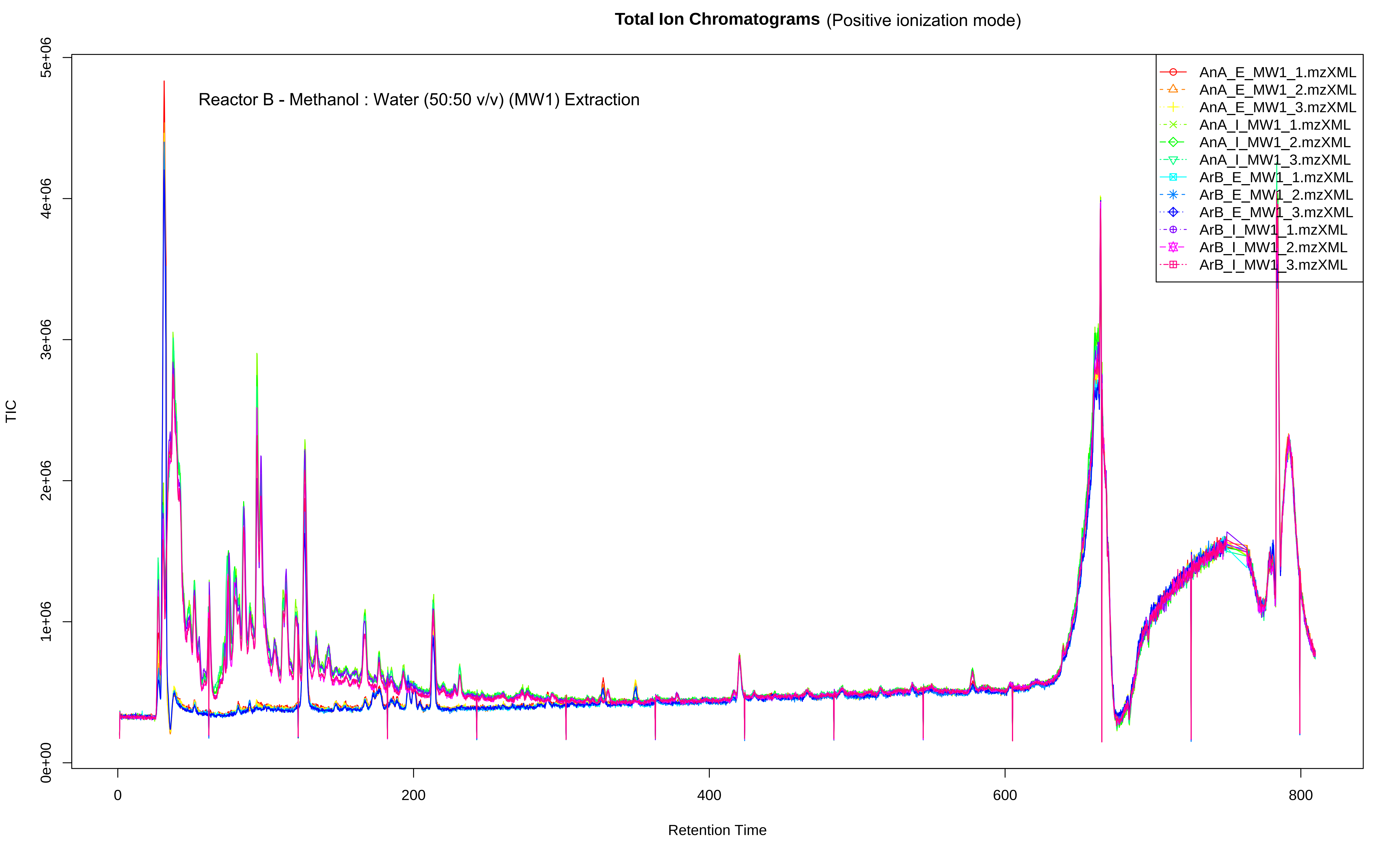

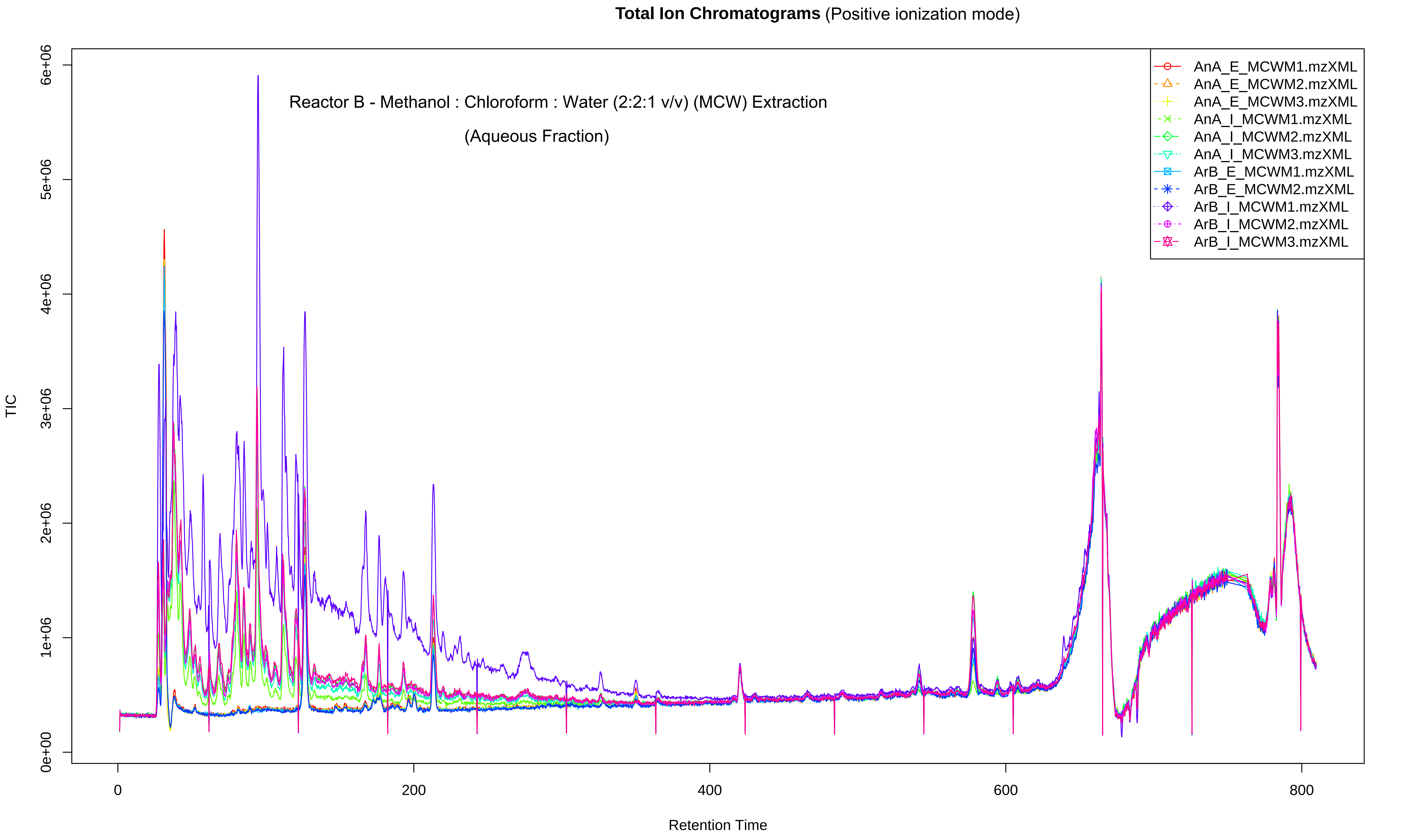

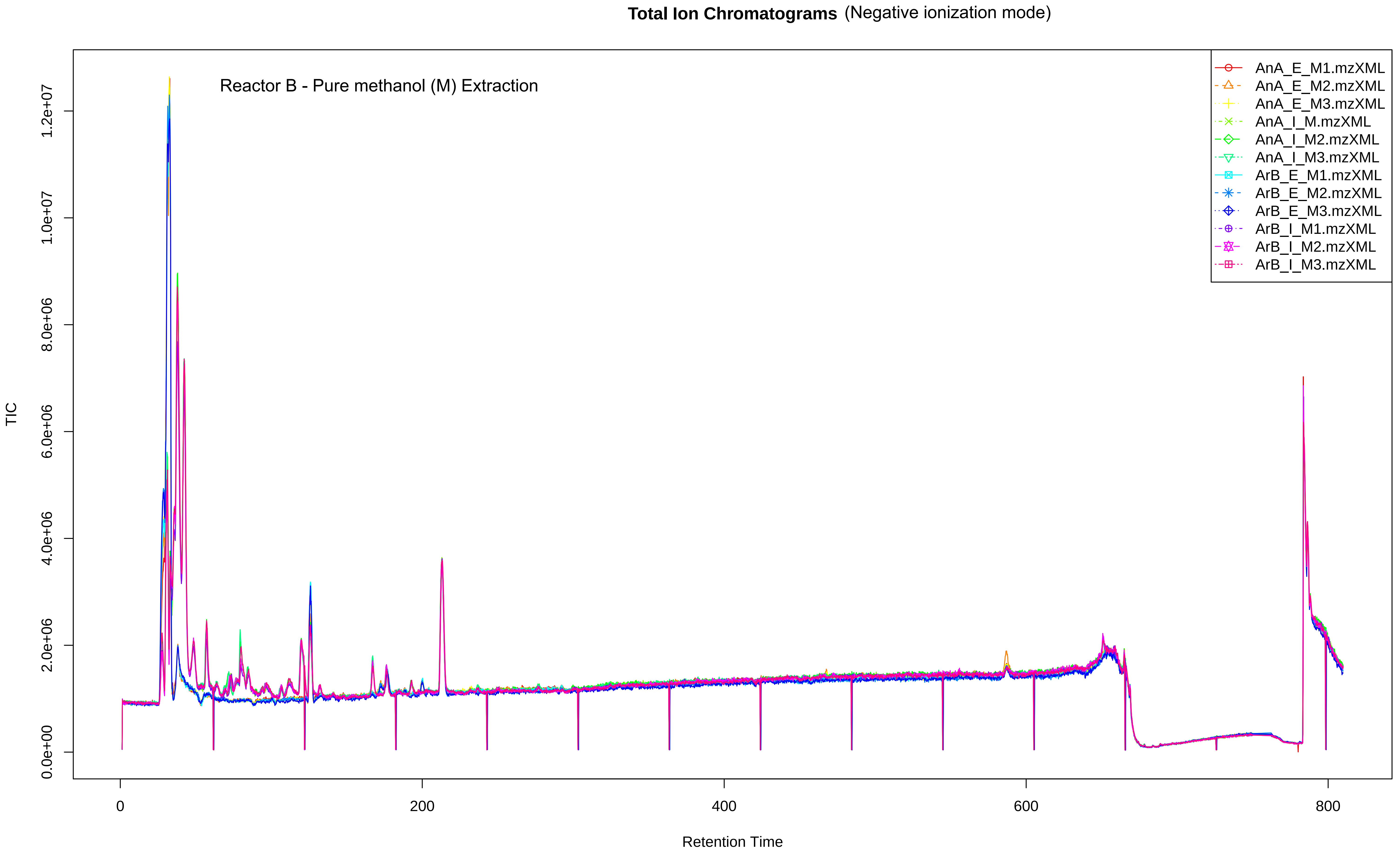

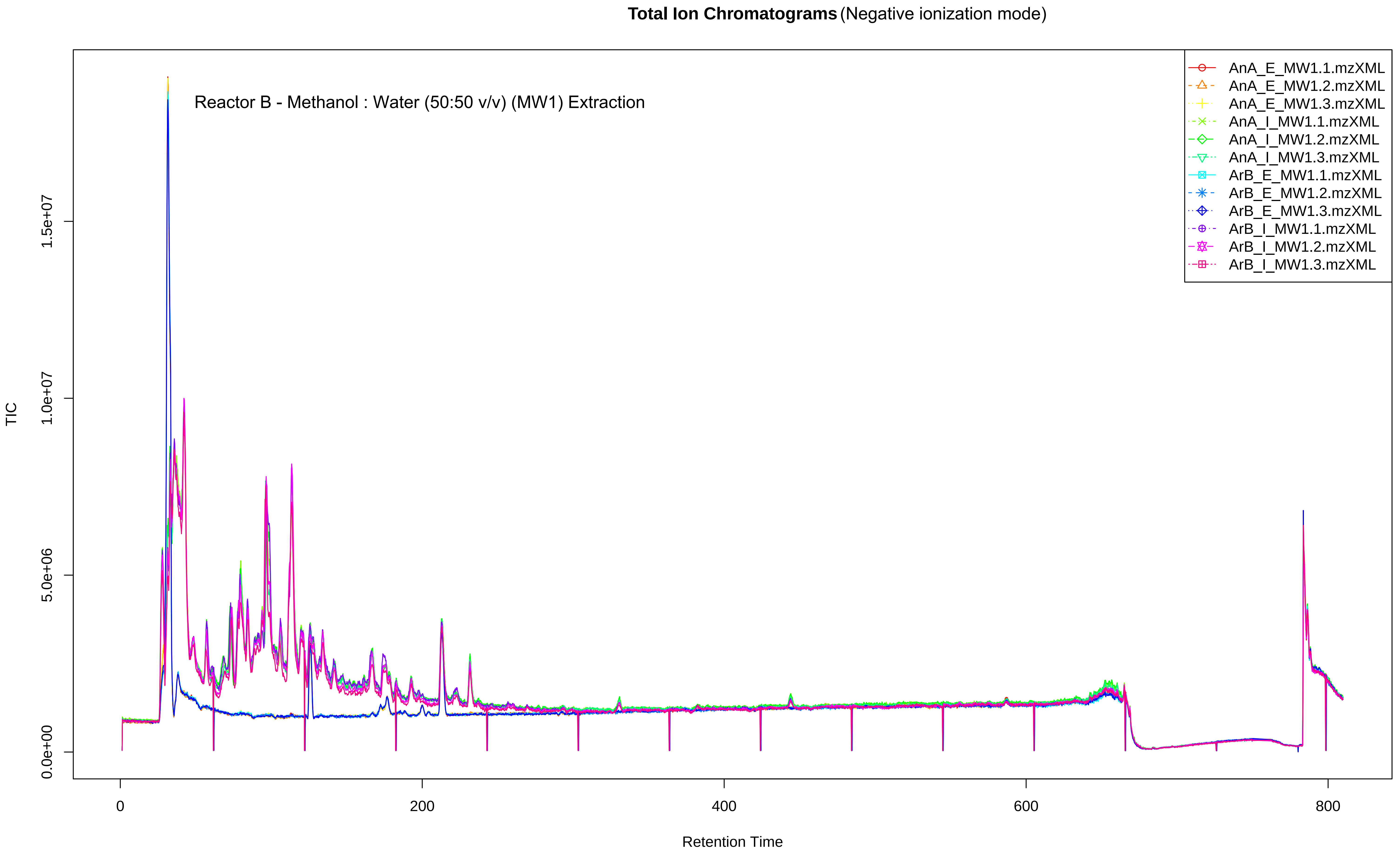

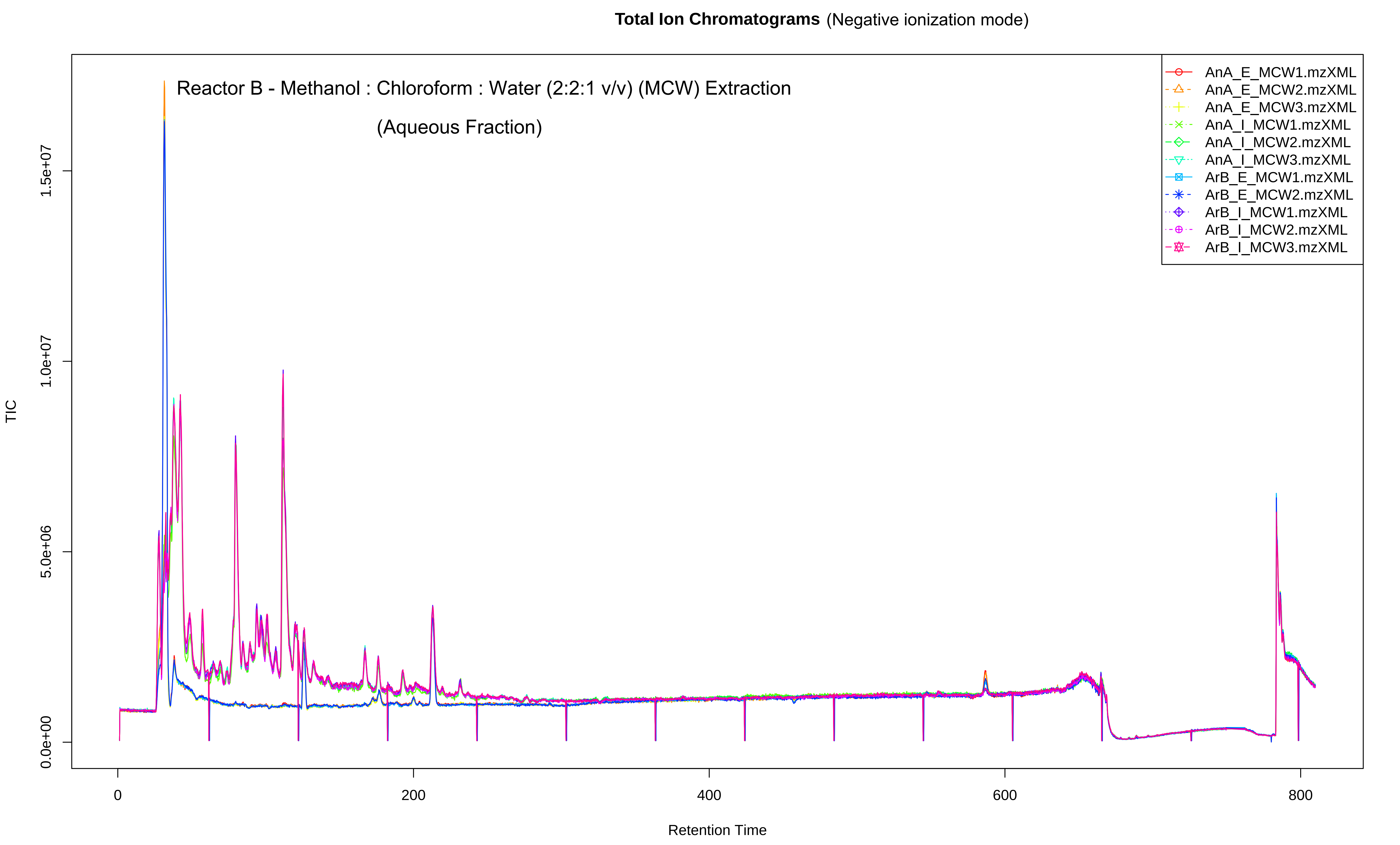

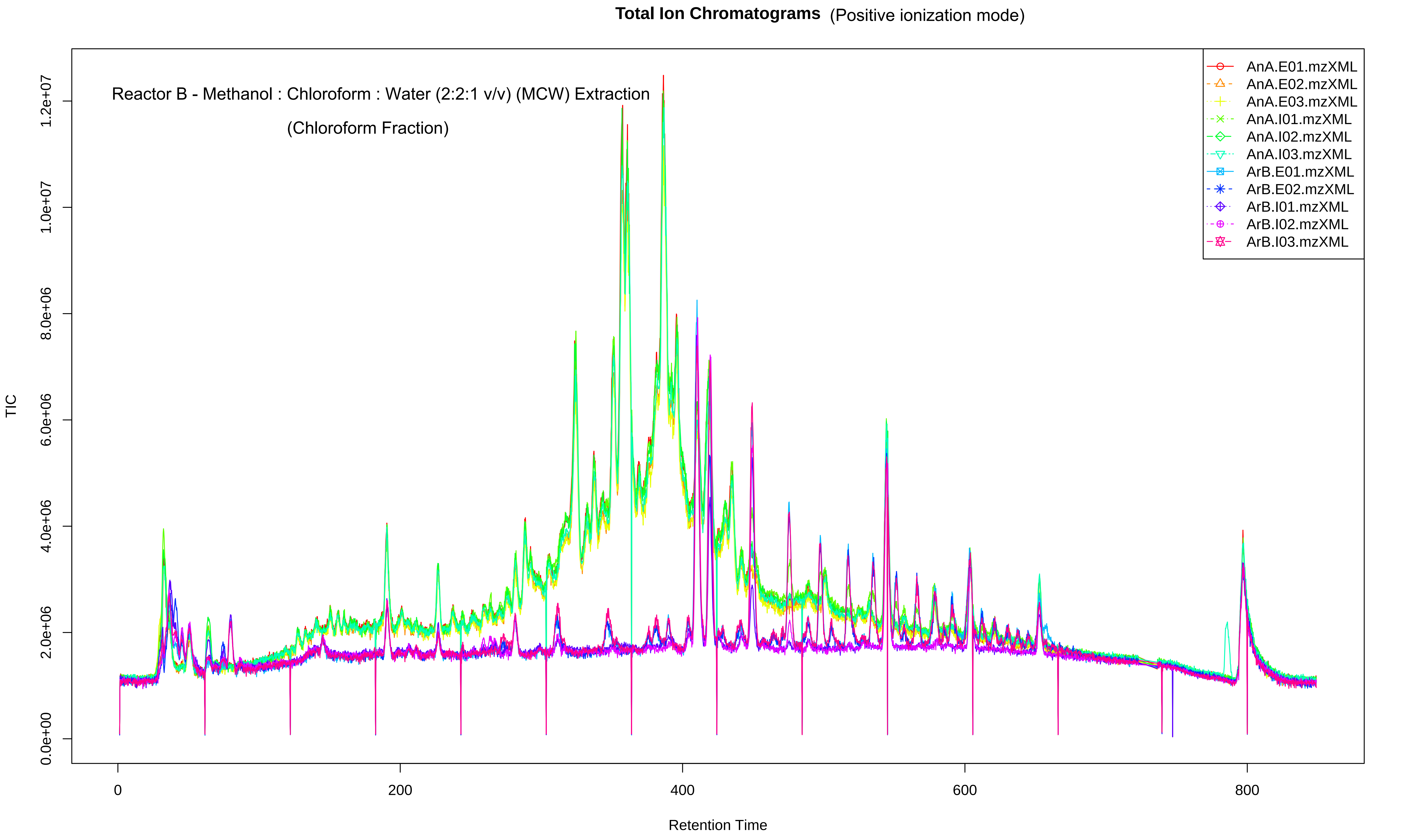

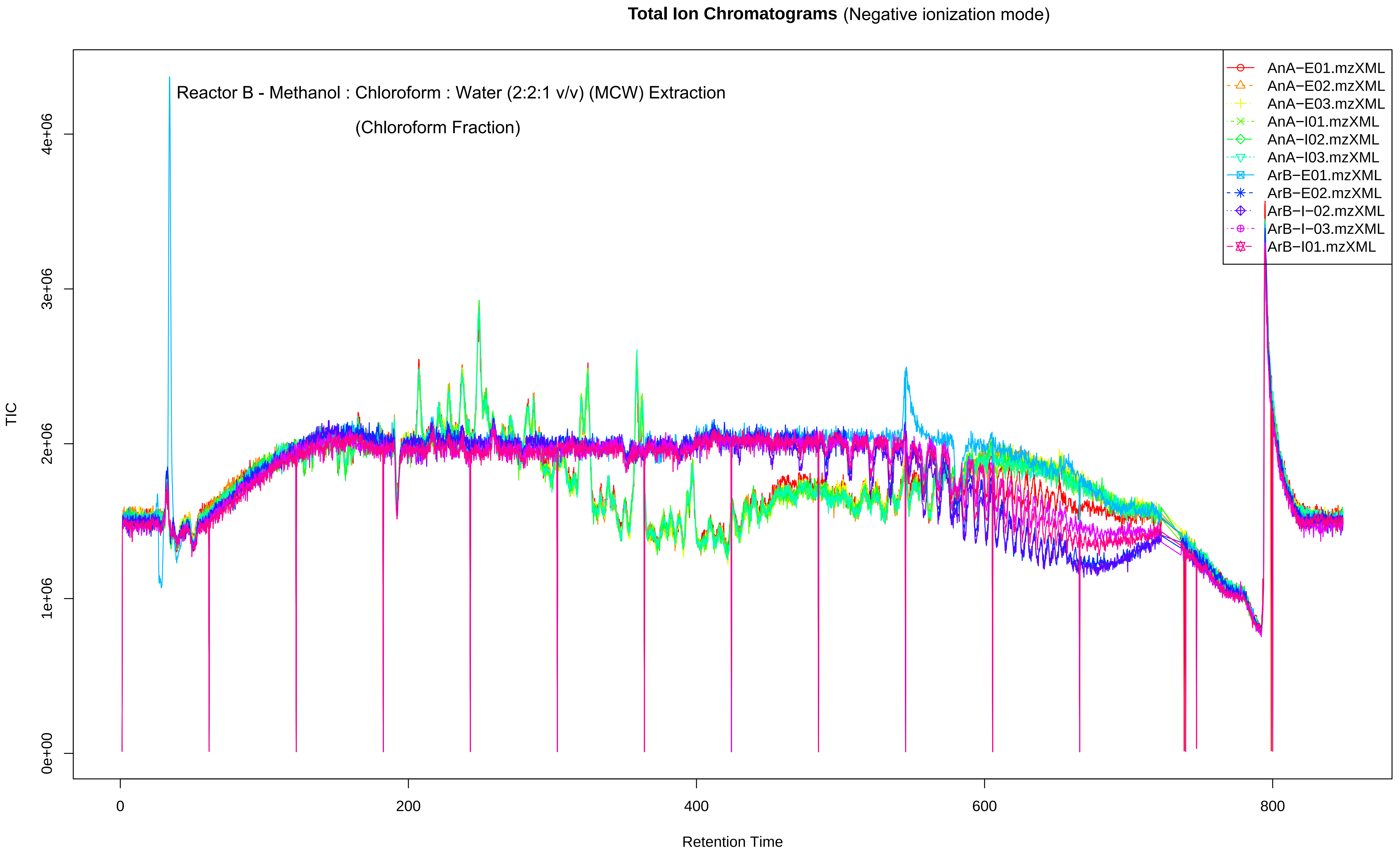
