## Supplementary Figures for "Evaluation of extraction solvents for untargeted metabolomics analysis of enrichment reactor cultures performing enhanced biological phosphorus removal (EBPR)"

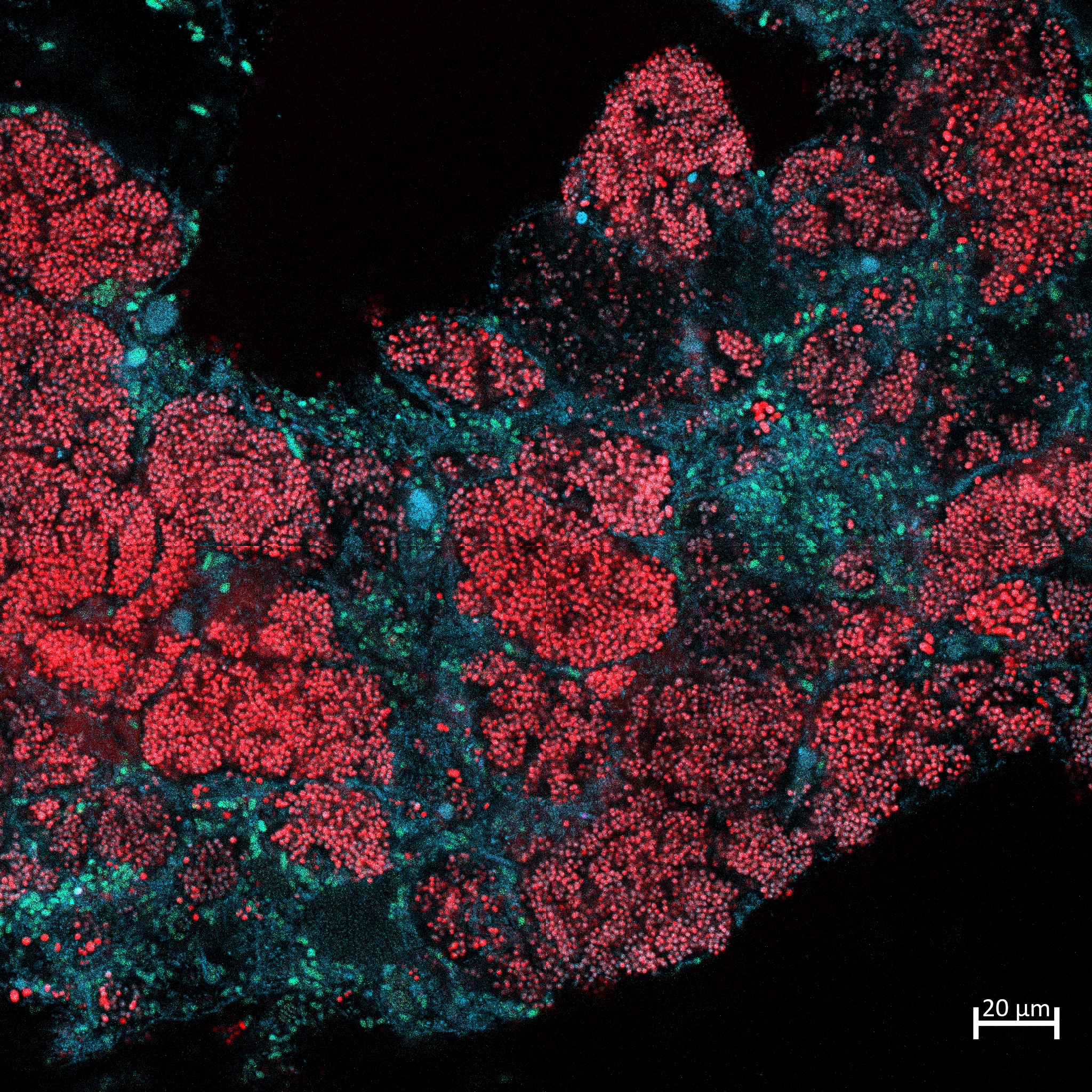

Fig.1 Fluorescence in situ hybridisation (FISH) composite image of biomass sampled from the enriched community in Reactor A. Biomass was labelled with probes PAOMIX (red; targeting *Ca.* Accumulibacter) and EUBMIX (green; universal bacterial probe).

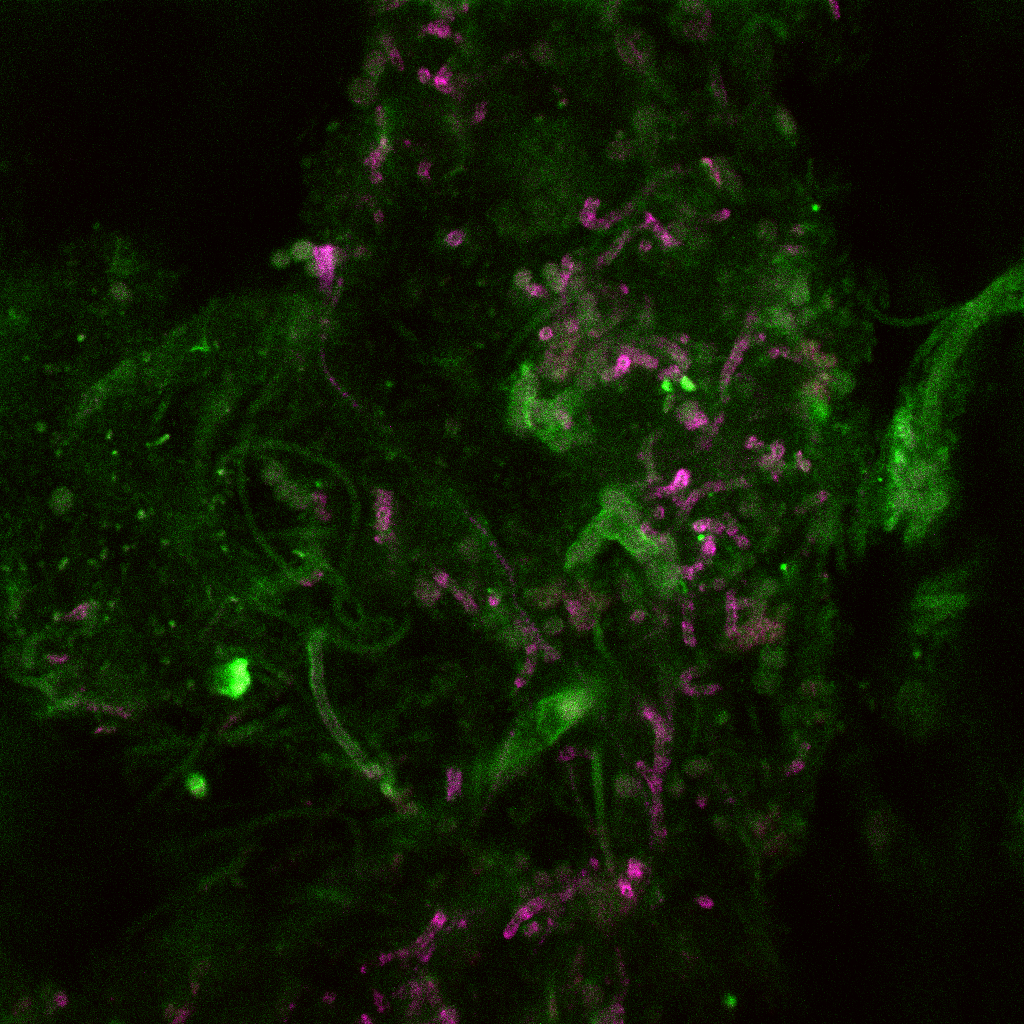

Fig. 2 Fluorescence in situ hybridisation (FISH) composite image of biomass sampled from the enriched community in Reactor B. Biomass was labelled with probes Actino-221 and Actino-658 (magenta; targeting *Tetrasphaera*), and EUBMIX (green; universal bacterial probe).

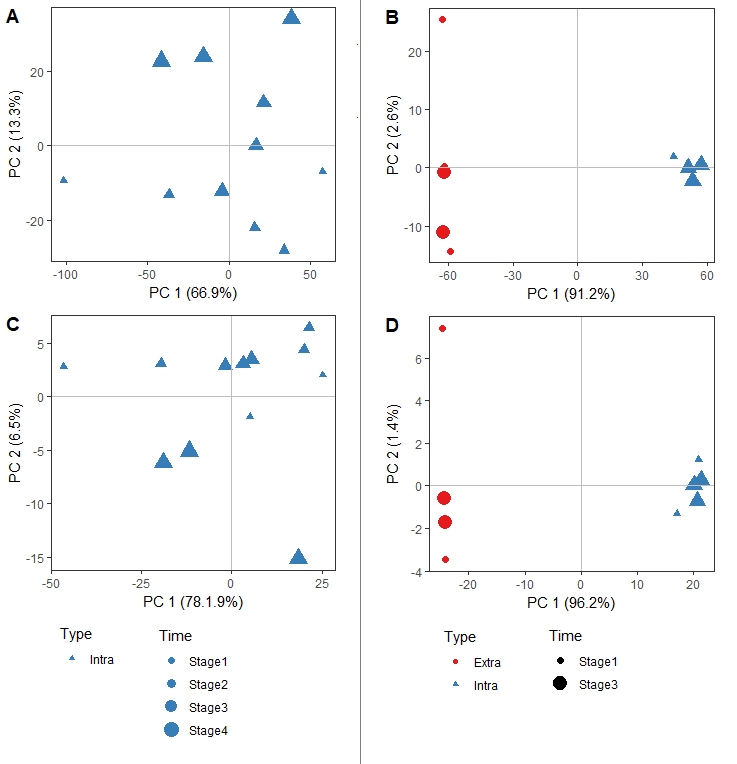

Fig 3 PCA analysis of non-polar metabolite profiles extracted by MCW from Reactor A (Panel A and C) and Reactor B (Panel B and D), and categorised by ionisation mode (positive mode in panels A-B; negative mode in panels C-D).

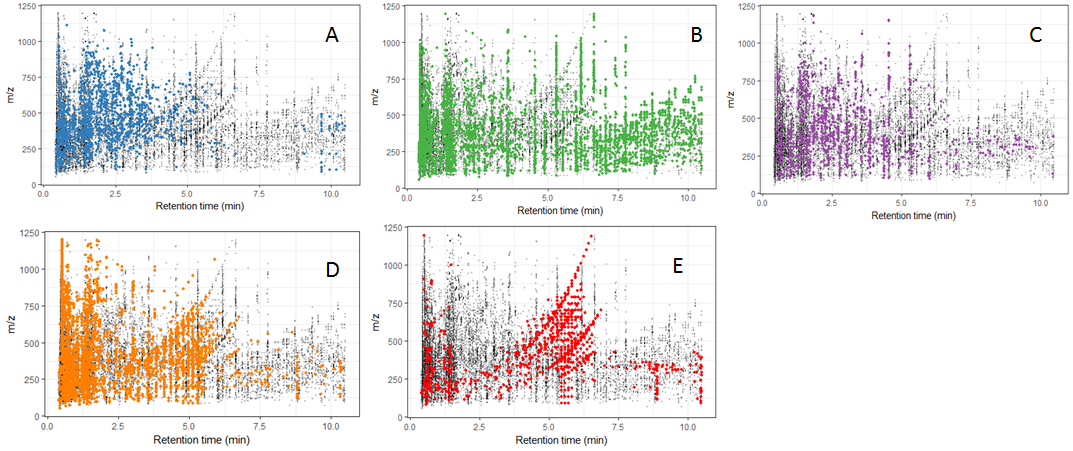

Fig. 4: Map of the feature positions in the rt/mz plane, positive ionization mode for Reactor A data. Gray points indicate features found in all the samples. Colored points show features in which the highest mean value is observed compared to other extraction methods: *A*) methanol:chloroform:water (40:20:20 v/v) (MCW), *B*) methanol:water (50:50) (MW1), *C*) methanol:water (60:40) (MW2), *D*) methanol:water (80:20) (MW3) and *E*) pure methanol (99.99%).

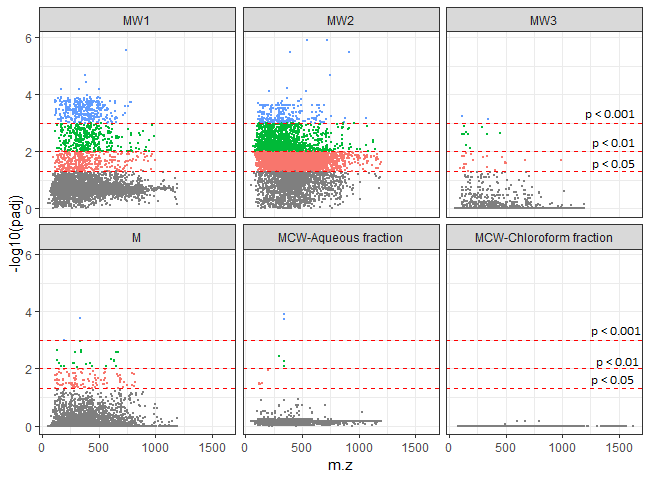

Fig. 5: Analysis of differential abundance of mass feature between different sampling points for each extraction methods in Reactor A. Horizontal-axis shows m/z ratio and vertical axis shows the adjusted P-value of ANOVA test statistic (time point as a single factor). Horizontal lines and colour coding indicate different adjusted P-value thresholds: P = 0.05, P = 0.01 and P = 0.001.

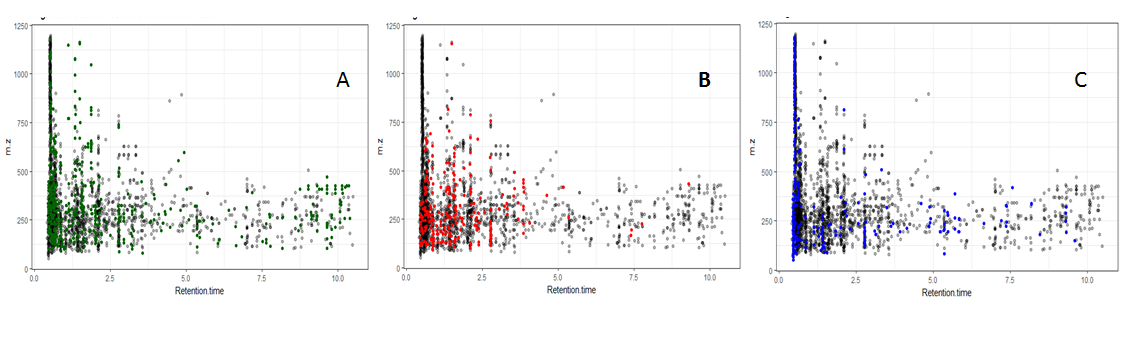

Fig. 6: Map of the feature positions in the rt/mz plane, positive ionization mode for Reactor B data. Gray points indicate features found in all the samples. Colored points show features in which the highest mean value is observed compared to other extraction methods: *A*) methanol:chloroform:water (40:20:20 v/v) (MCW), *B*) methanol:water (50:50) (MW1), *C*) pure methanol (99.99%).

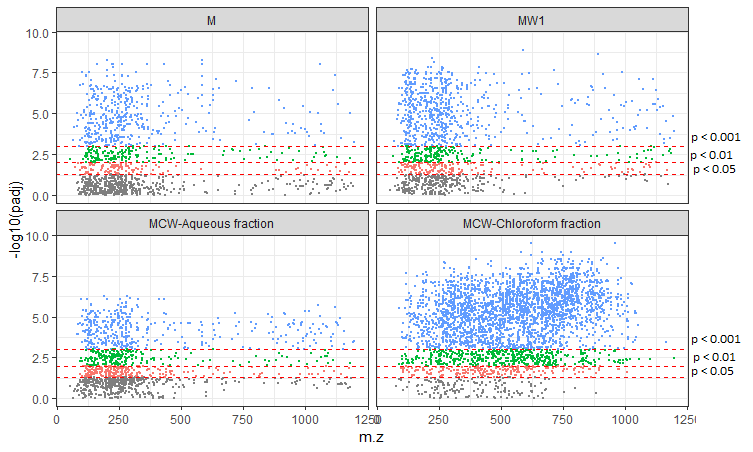

Fig. 7 Analysis of differential abundance of mass feature between different sampling points for each extraction methods in Reactor B. Horizontal-axis shows m/z ratio and vertical axis shows the adjusted P-value of ANOVA test statistic (four time point-compartment combiantions treated as a single factor). Horizontal lines and colour coding indicate different adjusted P-value thresholds: *P* = 0.05, *P* = 0.01 and *P* = 0.001.
